# Compartment-Specific Neurexin Nanodomains Orchestrate Tripartite Synapse Assembly

**DOI:** 10.1101/2020.08.21.262097

**Authors:** Justin H. Trotter, Zahra Dargaei, Alessandra Sclip, Sofia Essayan-Perez, Kif Liakath-Ali, Karthik Raju, Amber Nabet, Xinran Liu, Markus Wöhr, Thomas C. Südhof

**Author notes:** These authors contributed equally to the study (Justin H. Trotter, Zahra Dargaei). Laboratory for Behavioral Neuroscience, Department of Biology, Faculty of Science, University of Southern Denmark, Campusvej 55, DK-5230 Odense M, Denmark, and Behavioral Neuroscience, Experimental and Biological Psychology, Faculty of Psychology, Philipps-University of Marburg, Gutenbergstraße 18, D-35032 Marburg, Germany.

## Abstract

At tripartite synapses, astrocytes enmesh synaptic contacts, but how astrocytes contribute to the formation, maturation and plasticity of synapses remains elusive. Here we show that both astrocytes and neurons abundantly express neurexin-1, a presynaptic adhesion molecule that controls synaptic properties. Using super-resolution imaging, we demonstrate that presynaptic neuronal and astrocytic neurexin-1 form discrete nanoclusters at excitatory synapses. We find that distinct patterns of heparan sulfate modification and alternative splicing confer onto astrocytic and neuronal neurexin-1 different ligand specificities, thereby enabling compartment-specific signaling by neurexin-1. At hippocampal Schaffer-collateral synapses, deletion of neurexin-1 from either astrocytes or neurons did not alter synapse numbers, but differentially impaired synapse function. Neuronal neurexin-1 was essential for NMDA-receptor-mediated synaptic responses, whereas astrocytic neurexin-1 was required for maturation of silent synapses, AMPA-receptor recruitment, and long-term potentiation. Thus, astrocytes and neurons surprisingly use the same synaptic adhesion molecule to control distinct synapse properties.

---

Synapses in brain are often physically coupled to astrocytic processes, thus forming tripartite synapses^1–3^. Astrocytes contribute to numerous brain functions, including neurotransmitter metabolism, neurovascular coupling, synaptogenesis, and synaptic pruning^4–6^. In particular, astrocytes promote synapse formation through the secretion of synaptogenic factors (e.g. SPARCL1, thrombospondins, ApoE, Chordin-like 1, TGFβ, and glypicans)^7–12^ and via contact-dependent mechanisms^13–15^. However, the role of astrocytes in synaptogenesis and synaptic pruning does not readily explain why they make persistent intercellular junctions with synapses well after their formation and into adulthood. Rather, these junctions may serve as intercellular signaling hubs that enable bidirectional communication between astrocytes and neurons.

Diverse synaptic adhesion molecules guide developmental processes such as neuronal positioning, axonal targeting, synapse formation, and specification of synaptic properties^16, 17^. At least some of these synaptic adhesion molecules may also act in astrocytes to enable intercellular communication with neurons at tripartite synapses. To explore this possibility, we performed an expression screen of major synaptic adhesion molecules in astrocytes. We quantified the levels of actively translated mRNAs in astrocytes by crossing Cre-dependent ‘RiboTag’ mice with Cre-driver mouse lines that express Cre only in astrocytes (Aldh1l1-CreERT2^18^, induced with tamoxifen). Using highly purified ribosome-bound mRNAs from astrocytes (Extended Data Fig. 1a), we detected numerous synaptic adhesion molecules that were modestly (0.5 to 1-fold) to highly enriched (>1-fold) in astrocytes compared to total RNA (Fig. 1a). Interestingly, the risk for autism spectrum disorder (ASD) was prominently associated with synaptic adhesion genes exhibiting a 0.5-fold or greater enrichment in astrocytes. Most notably, we found that the synapse organizer neurexin-1 (Nrxn1) was highly enriched in astrocytes. Copy number variations that selectively alter expression of the human *NRXN1* gene are among the more frequent single-gene mutations observed in patients with ASD, schizophrenia, Tourette syndrome, and other neurodevelopmental disorders^19, 20^, suggesting that heterozygous loss-of-function of *NRXN1* may predispose to neuropsychiatric diseases in part by altering astrocytic functions.

**Figure 1.**
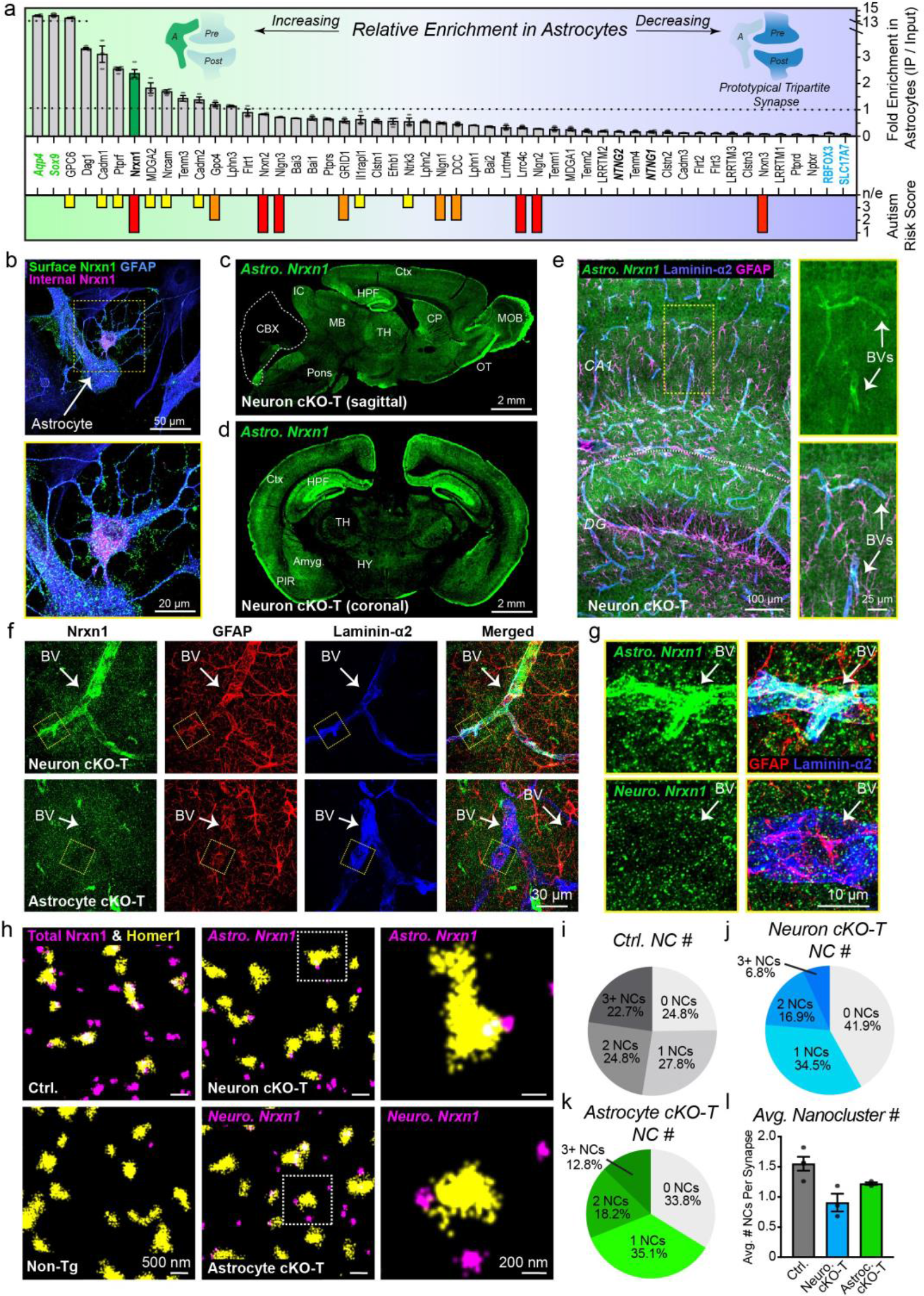
Astrocytes express and target the major synapse organizer Nrxn1 to vascular endfeet and synapses. **a**, Profile of synapse adhesion molecule enrichment in ribosome-bound mRNAs from hippocampal astrocytes compared to total mRNA (input) following quantitative RT-PCR (top). Astrocyte control genes (left, bold green) indicate the upper limit of enrichment, and neuron control genes the lower limit (right, bold blue). The SFARI autism risk score (1-3; found at gene.sfari.org) for each gene is denoted (bottom) including 1 (high confidence), 2 (strong candidate), and 3 (suggestive evidence). Bold and italicized genes are associated with syndromic autism. **b,** Surface and internal Nrxn1 (green and red, respectively) visualized in cultured glia by immunocytochemistry for HA and GFAP, an astrocyte marker. **c-d**, Selective visualization of astrocytic Nrxn1 in sagittal (**c**) and coronal (**d**) sections from Nrxn1 HA-knockin mice to pan-neuronal Cre mice to induce neuron-specific conditional knockout truncation (cKO-T). **e,** Representative image in the CA1 and dentate gyrus (DG) of astrocytic Nrxn1 (green) staining in neuron cKO-T mice co-labelled with GFAP (astrocytes, magenta) and laminin-alpha2 (blood vessels or BVs, blue). Yellow box, enlarged on right. **f-g**, Nrxn1 enrichment at astrocytic endfeet (GFAP, red) that target blood vessels (laminin-α2, blue) is detectable in the CA1 region of Nrxn1 neuron cKO-T (top) but not Nrxn1 astrocyte cKO-T mice (bottom). Yellow inset box enlarged on right (**g**). **h,** Representative images of Nrxn1 nanoclusters (purple) and Homer1 (yellow) visualized using direct stochastic optical reconstruction microscopy. Brain sections stained include HA-Nrxn1 control (Ctrl, top left), non-transgenic (Non-Tg, bottom left), Nrxn1 neuron cKO-T (top middle), and Nrxn1 astrocyte cKO-T (bottom middle; white inset boxes, magnified on right). Signal remaining following loss of neuronal Nrxn1 is considered to be astrocytic Nrxn1, whereas signal remaining after loss of astrocytic Nrxn1 is considered to be neuronal. **i-k**, Pie charts showing the distribution of Homer-positive excitatory synapses without or with Nrxn1 nanoclusters (i.e. 1, 2, and 3+) in HA-Nrxn1 ctrl (**i**), neuron cKO-T (**j**), and astrocyte cKO-T mice (**k**). **l**, Summary graph of the average number of nanoclusters per excitatory synapse. Data in **a** and **l** are mean ± SEM. *n =* 3-4 independent biological replicates per gene (**a**); 198 synapses / 4 mice (**i**), 148 synapses / 3 mice each (**j, k**); avg. per mouse for 4 Ctrl. and 3 per neuron & astrocyte cKO-Ts (**l**). *Abbreviations*: Ctx, cortex; IC, inferior colliculus; HPF, hippocampal formation; CBX, cerebellar cortex; MB, midbrain; TH, thalamus; CP, caudate putamen; OT, olfactory tract; MOB, main olfactory bulb; TH, thalamus; PIR, piriform cortex; HY, hypothalamus.

Neurexins are encoded by three genes (*Nrxn1*-*3* in mice) that contain independent promoters to drive transcription of long α-neurexins, shorter β-neurexins, and a very short neurexin-1 γ−isoform^21–25^. Extracellularly, α-neurexins possess six laminin/neurexin/sex hormone–binding globulin (LNS) domains that are interspersed with EGF-like repeats. In contrast, β-neurexins contain a short β-specific N-terminal sequence that splices into the α-neurexin sequence upstream of their sixth LNS domain. Following the sixth LNS domain, neurexins contain a heavily glycosylated “stalk” region that includes a site for heparan sulfate (HS) modification^26^, a cysteine-loop domain, a transmembrane region, and a cytoplasmic tail (Fig. 2a). Neurexin mRNAs are extensively alternatively spliced at six canonical homologous sites producing thousands of isoforms^27–29^ that are differentially expressed by neuronal subclasses^30, 31^. Presynaptic neurexins concentrate in nanoclusters^32^, and interact trans-synaptically, often in a manner modulated by alternative splicing, with a multitude of postsynaptic ligands^20, 33–44^. Via their trans-synaptic interactions, neurexins specify diverse synaptic properties, including presynaptic Ca^2+^ channels^45–,47^, tonic postsynaptic endocannabinoid synthesis^48^, GABAB-receptors^49^, BK potassium channels^47^ and trans-synaptic recruitment of postsynaptic α-amino-3-hydroxy-5-methyl-4-isoxazolepropionic acid receptors (AMPARs)^50^ and N-methyl-D-arginine–type glutamate receptors (NMDARs)^51^. The finding that Nrxn1 is highly expressed in astrocytes (Fig. 1a) suggests that it may perform its synaptic functions, at least in part, via expression in astrocytes, thereby enabling astrocytes to signal through synaptic ligands and to regulate diverse synaptic properties.

**Figure 2.**
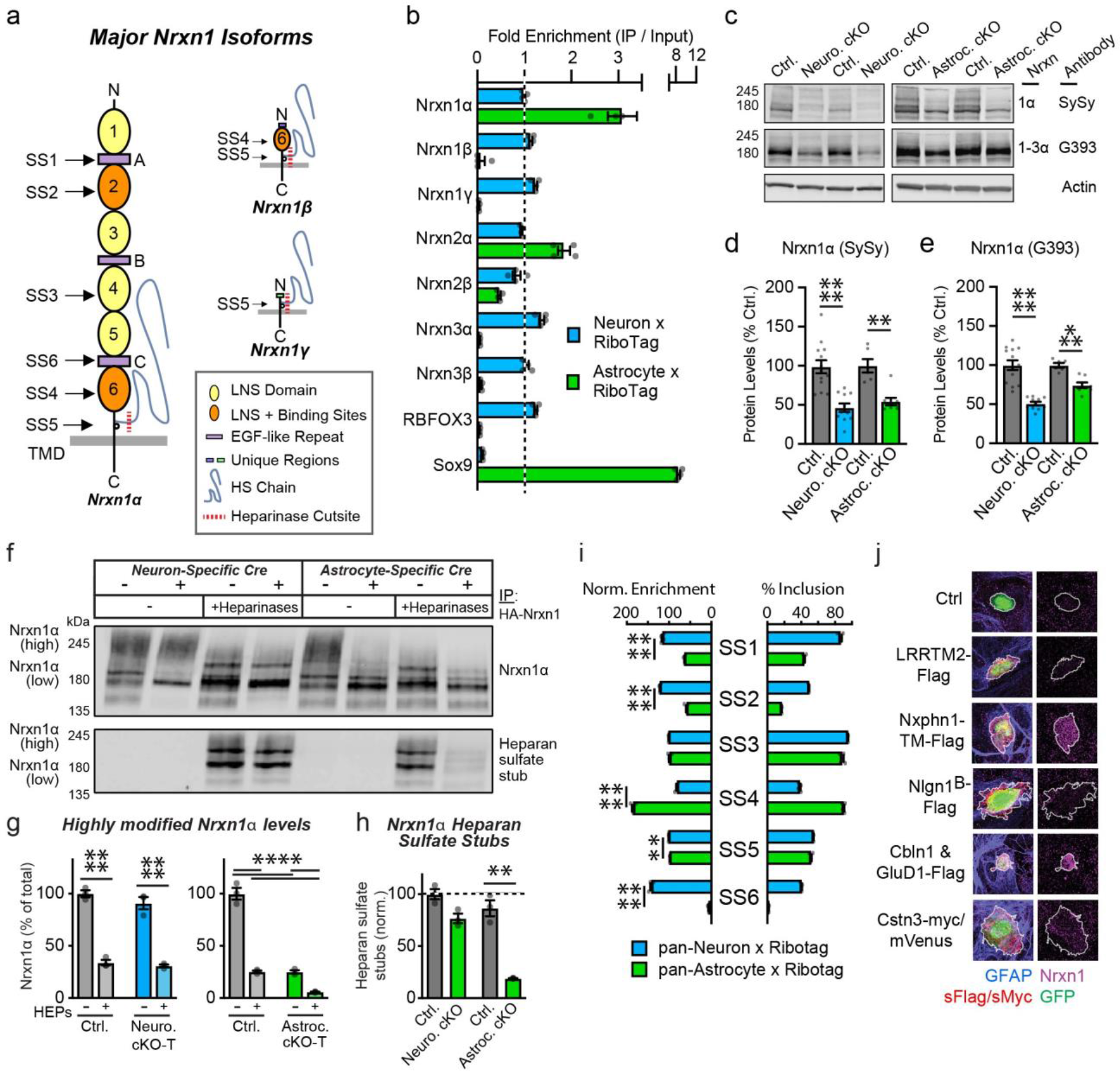
Astrocytes and neurons employ distinct molecular programs that involve Nrxn1. **a,** Cartoon of major Nrxn1 isoforms (i.e., α, β, and γ). The location of splice sites 1-6 (SS1-SS6) are indicated to the left. Bottom right, key of major structural features. **b,** Quantifications of major neurexin isoform transcripts via quantitative RT-PCR using either astrocyte- or neuron-specific ribosome-bound mRNAs (targeted using ‘RiboTag’) and compared to total RNA (input). **c,** Representative immunoblots of Nrxn1α or Nrxn1/2/3α using antibodies with different degrees of specificity. Total brain lysates from neuron- and astrocyte-specific Nrxn1 cKO mice and littermate controls were examined. **d-e**, Summary graphs of Nrxn1α (**d**) or pan-Nrxn1/2/3α (**e**) levels normalized to beta-actin and expressed as percent Ctrl levels. **f-h,** Representative immunoblots (**f**) of Nrxn1α in hippocampal lysates from Nrxn1 neuron and astrocyte cKO-T mice (cre+) and littermate ctrls. Nrxn1 was immunoprecipitated followed by immunoblotting without or with prior heparinase treatment. Summary graphs of highly-modified Nrxn1α, which is equivalent to heparan sulfate-modified Nrxn1α (**g**). Heparinase treatment of immunoprecipitated Nrxn1 from brain lysates almost completely converts highly modified Nrxn1α into its less modified form. Relative levels of heparan-sulfate stubs, normalized for total Nrxn1α levels (**h**). **i,** Quantification of Nrxn1 alternative splicing at splice sites 1-6 (SS1-SS6) in either neuron or astrocyte-bound mRNA purified using the RiboTag approach following by junction-flanking RT-PCR. **j,** Representative images of surface Nrxn1 (magenta) recruitment by a subset of Nrxn1 ligands (red) expressed on the surface of HEK293T cells (GFP, green) and co-cultured with HA-Nrxn1 cKI primary astrocytes (GFAP, blue). Numerical data are means ± SEM. *n =* 4 mice for neuron mRNAs and 8 mice with 2 mice pooled per pull-down for astrocyte mRNAs (**b, i**); 11 neuron cKO and 13 littermate ctrls., 8 astrocyte cKO and 6 littermate ctrls. (**d-e**); 3 mice per group (**f-h**). Statistical significance was assessed with a two-way ANOVA with a Tukey’s post-hoc test (**g**) or two-tailed unpaired t-test to controls (rest), with ** = p < 0.01; *** = p < 0.001; **** = p < 0.0001.

## Astrocytic Nrxn1 forms contacts with both neurons and blood vessels

We confirmed by single-molecule RNA *in situ* hybridization that *Nrxn1* expression is abundant in both neurons and astrocytes throughout the hippocampus (Extended Data Fig. 1b-c). Using astrocytes cultured from HA-tagged Nrxn1 conditional knockin (cKI) mice that permit highly specific localization of endogenous Nrxn1 with HA antibodies^32^, we found that Nrxn1 was present on the surface of astrocytes in a largely punctate pattern, consistent with a localization to specific plasma membrane domains (Fig. 1b). We then selectively visualized astrocytic Nrxn1 in the brain by crossing HA-Nrxn1 cKI mice to a pan-neuronal Cre-driver line^52^ that was validated with a reporter mouse line (Extended Data Fig. 1d). Cre-recombination of the HA-Nrxn1 cKI locus using the pan-neuronal Cre-driver line produces an inactive truncated Nrxn1 protein (cKO-T) in neurons (Extended Data Fig. 4g)^32^. Since the truncated HA-Nrxn1 after Cre-recombination is rapidly degraded and only faintly detectable by immunocytochemistry, any HA-Nrxn1 detected after neuron-specific Cre-recombination of HA-Nrxn1 cKI mice is from astrocytes. In brain sections, we observed widespread but highly heterogeneous expression of astrocytic Nrxn1 including in the hippocampus, the cortex, the olfactory tract and olfactory glomeruli (Fig. 1c-d). In contrast, astrocytic Nrxn1 was expressed sparsely in the midbrain and virtually absent from the cerebellum. The remarkable diversity of astrocytic Nrxn1 expression patterns observed across the brain is consistent with recent reports of the circuit-specific functional and molecular diversity of astrocytes^53–55^.

To directly compare the levels and distribution of neuronal and astrocytic Nrxn1, we also crossed the HA-Nrxn1 cKI mice with Aldh1l1-CreERT2 Cre-driver mice, which enable the tamoxifen-inducible truncation of HA-Nrxn1 in astrocytes. Astrocyte-specific expression of Cre in this mouse model was validated using a reporter mouse line (Extended Data Fig. 1e). Imaging revealed that neurons and astrocytes separately contribute approximately half of total Nrxn1 levels in the hippocampal neuropil, with both cellular pools exhibiting subfield-specific enrichments (Extended Data Fig. 2a-c). Strikingly, we found that much of the astrocytic Nrxn1 was trafficked to vascular endfeet, which encapsulate blood vessels (Fig. 1e; Extended Data Fig. 3a). Confirming the astrocytic origin of the vascular labeling, HA-Nrxn1 labeling of astrocytic endfeet was specific, and was abolished in Nrxn1 astrocyte cKO tissue (Fig 1f-g; Extended Data Fig. 3a). Collectively, these results reveal that astrocytes abundantly express Nrxn1 and transport it to two separate target domains, the synaptic neuropil and vascular endfeet.

**Figure 3.**
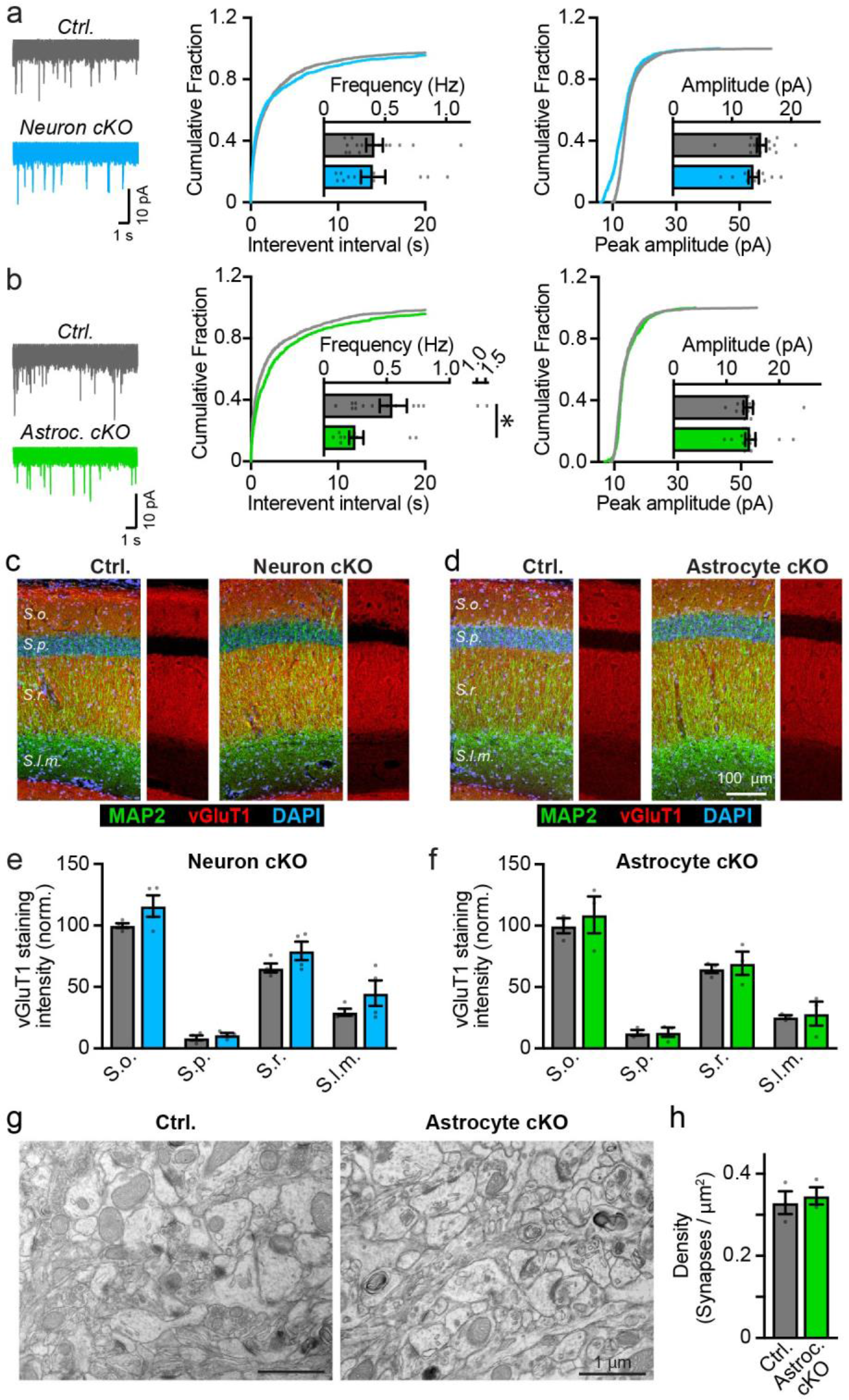
Deletion of Nrxn1 from astrocytes suppresses spontaneous excitatory synaptic transmission without altering synapse numbers. **a-b,** Astrocyte-but not neuron-specific deletion of Nrxn1 in cKO mice reduces the mEPSC frequency without changing the mEPSC amplitude. mEPSCs were recorded at P22-P26 in acute slices from CA1 pyramidal neurons prepared from neuron- (**a**), and astrocyte-specific Nrxn1 cKOs (**b**). Representative traces are shown (left), as well as summary graphs of the mEPSC frequency and cumulative plots of interevent interval (middle), and summary graphs and cumulative plots of amplitude (right). **c-d,** Representative sections of the hippocampal CA1 region from control mice and mice with neuron- (**c**) or astrocyte-specific deletion of Nrxn1 (**d**) were stained for the excitatory presynaptic marker vGluT1 (red), the dendritic marker MAP2, and the nuclear marker DAPI (blue). Smaller panels to the right of main panels show only vGluT1 staining as a proxy for synapse density (*S.o*., *Stratum oriens*; *S.p*., *Stratum pyramidale*; *S.r.*, *Stratum radiatum*; *S.l.m.*, *Stratum lacunosum-moleculare*). **e-f,** Summary graphs quantify the vGluT1 staining intensity (normalized to that of MAP2 and expressed as % of *Stratum oriens* levels) for the indicated layers. **g-h,** Electron microscopy reveals that deletion of Nrxn1 from astrocytes does not alter excitatory synapse density in the *Stratum radiatum* of the CA1 region (**g**, representative electron micrographs of the neuropil; **h**, summary graph of synapse density). Numerical data are means ± SEM. *n =* 15 cells / 4 mice ctrl., 11 cells / 4 mice neuron cKO (**a**); 14 cells / 4 mice ctrl., 13 cells / 3 mice astrocyte cKO (**b**); 4 mice both ctrl. and neuron cKO (**e**); 3 mice both ctrl. and astrocyte cKO (**f, h**). Statistical significance was determined by two-tailed Mann Whitney test (**a**, **b**) or two-tailed unpaired t-test to controls (**e**, **f**, **h**), with * = p < 0.05; ** = p < 0.01.

**Figure 4.**
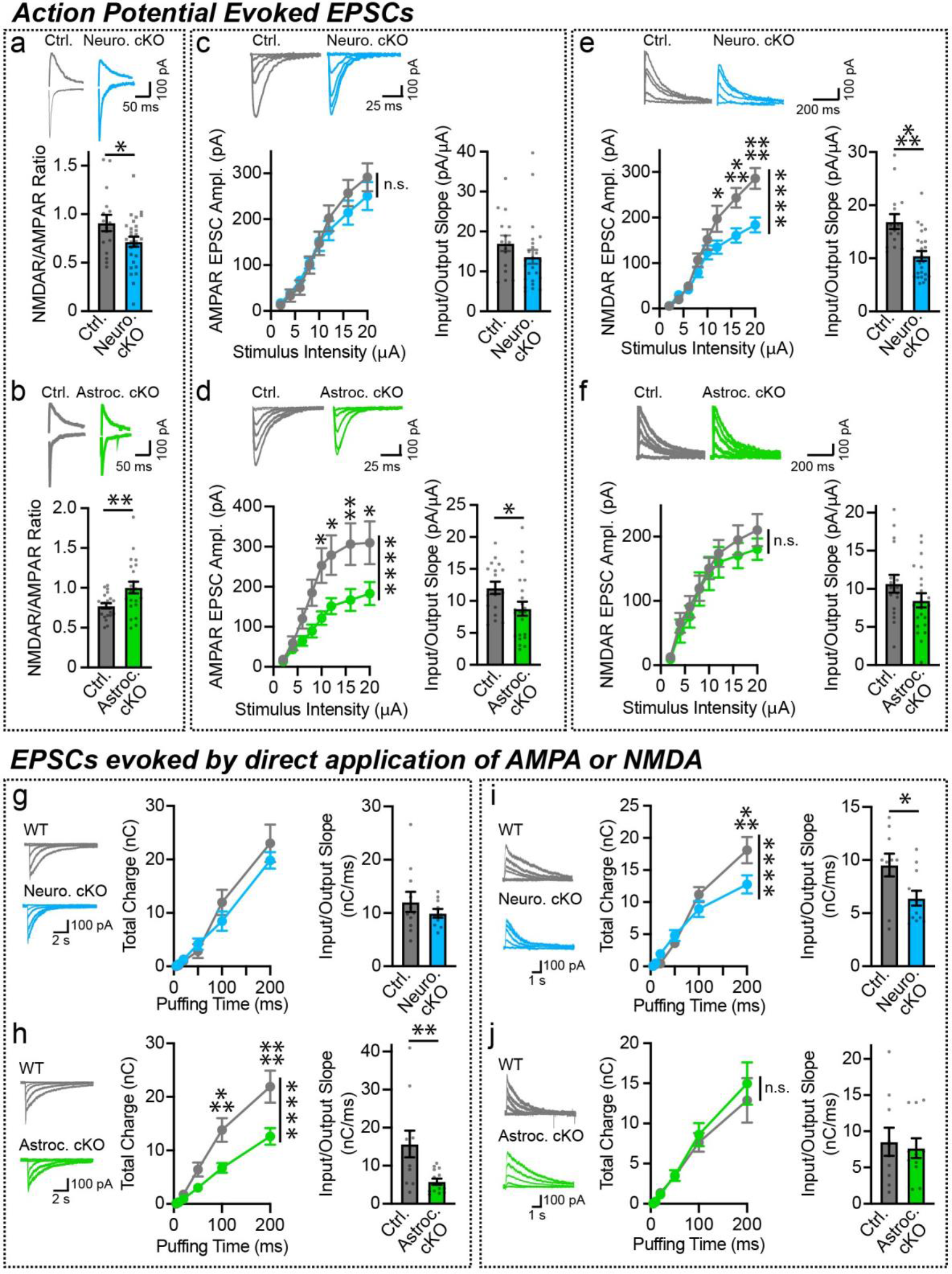
Neuronal and astrocytic Nrxn1 regulate distinct postsynaptic functions. **a-f,** EPSCs were evoked by electrical stimulation of the Schaffer collaterals, and recorded from hippocampal CA1 pyramidal neurons in acute slices from P22-P26 mice of the indicated genotypes. Sample traces are shown on top; for C-F, input-output curves (left) and slope summary graphs (right) are shown separately. Neuron-specific deletion of Nrxn1 lowers the NMDAR/AMPAR ratio (**a**), whereas astrocyte-specific deletion of Nrxn1 increases the NMDAR/AMPAR ratio (**b**). Deletion of Nrxn1 in neurons has no effect on AMPAR-EPSC strength (**c)**, unlike deletion of Nrxn1 in astrocytes which suppresses AMPAR-EPSCs **(d**). In contrast, deletion of neuronal Nrxn1 decreases the strength of evoked NMDAR-EPSCs (**e**), but astrocytic Nrxn1 deletion has no effect (**f**). **g-j**, EPSCs were elicited by direct application of AMPA or NMDA agonists. Sample traces (left), charge transfer over time curves (middle), and slope summary graphs (right) are shown. Deletion of neuronal Nrxn1 does not alter postsynaptic currents elicited by AMPA puffing (**g**), whereas deletion of astrocytic Nrxn1 significantly reduces AMPA-elicited EPSCs (**h**). On the contrary, neuron-**(i**) but not astrocyte-specific deletion of Nrxn1 suppresses currents following application of NMDA. All data are means ± SEM. *n =* 16 cells / 5 mice ctrl., 28 cells / 5 mice neuron cKO (**a**), 21 cells / 5 mice ctrl., 22 cells / 4 mice astrocyte cKO **(b**); 15 cells / 4 mice ctrl., 24 cells / 5 mice neuron cKO (**c**, **e**); 17 cells, 5 mice ctrl., 22 cells / 6 mice astrocyte cKO (**d**, **f**); 11 cells / 3 ctrl., 10 cells / 3 neuron cKO (**g**); 11 cells / 3 ctrl., 14 cells / 4 astrocyte cKO (**h**); 10 cells / 4 ctrl., 12 cells / 4 neuron cKO (**i**); 10 cells / 3 ctrl., 11 cells / 3 astrocyte cKO (**j**). Statistical analysis was assessed by two-way ANOVA with Sidak’s multiple comparison test (all input-output and charge transfer plots), two-tailed unpaired t-test (**a**, **d**, and all input-out-output slopes), with * = p < 0.05; ** = p < 0.01; *** = p < 0.001; **** = p < 0.0001.

## Neuronal and astrocytic Nrxn1 assemble into distinct nanoclusters in tripartite synapses

Using direct stochastic optical reconstruction microscopy (dSTORM), we compared the nanoscale distribution of presynaptic neuronal and astrocytic Nrxn1 in the *Stratum radiatum* of the CA1 region, which receives Schaffer collateral (SC) inputs from CA3 neurons. Consistent with previous results, Nrxn1 nanoclusters were frequently associated with excitatory synapses that were visualized by staining the postsynaptic scaffold protein Homer1^32^. After loss of presynaptic Nrxn1 clusters upon truncation of HA-Nrxn1 using neuron-specific Cre expression, a sizeable synaptic population of astrocytic Nrxn1 nanoclusters remained that were associated with more than half of excitatory synapses. This observation is consistent with electron microscopy studies suggesting that ∼60% of SC-CA1 synapses are contacted by astrocytes^1, 56^. Deletion of either Nrxn1 in neurons or in astrocytes decreased the average number of Nrxn1 nanoclusters per synapse (Fig. 1h-k). Astrocytic Nrxn1 nanoclusters were significantly larger and contained more localization signals than presynaptic Nrxn1 nanoclusters (Extended Data Fig. 3b-e), suggesting that astrocytic and presynaptic neuronal Nrxn1 nanodomains are molecularly and functionally distinct.

## Distinct molecular programs compartmentalize Nrxn1 signaling within tripartite synapses

The existence of astrocytic and presynaptic Nrxn1 nanoclusters that target the same synapse presents a conundrum, given that both clusters may engage similar postsynaptic ligands. For example, how is signaling by these nanoclusters to synaptic ligands segregated in order to maintain the correct information flow at separate compartments within tripartite synapses (i.e., astrocyte-synapse and pre-/postsynaptic junctions)? Amongst the synapse organizers that are translated at moderate to high levels by astrocytes (Fig. 1a), Nrxn1 is unique because of its extensive alternative splicing that generates hundreds of isoforms^27–29^ and because it interacts with scores of secreted and postsynaptic ligands^20^. This prompted us to ask whether astrocytes and neurons achieve compartment-specific signaling by employing distinct molecular programs involving Nrxn1.

Quantification of major *Nrxn* isoforms using ribosome-bound mRNA purified from neurons or astrocytes (Extended Data Fig. 1a), revealed that while *Nrxn1*α and *Nrxn2*α mRNAs were abundant in both astrocytes and neurons, other neurexin mRNAs were enriched in neurons but de-enriched in astrocytes (Fig. 2b). To directly measure the amounts of Nrxn1 protein in astrocytes or neurons, we crossed *Nrxn1* conditional knockout (cKO) mice^32, 45^ with the neuron- and astrocyte-specific Cre-driver mice described above, resulting in mice with neuron- or astrocyte-specific deletions of *Nrxn1* (referred to, respectively, as ‘neuron cKO’ and ‘astrocyte cKO’ mice; Extended Data Fig. 4a-d). Astrocyte and neuron *Nrxn1* cKO mice survived normally into adulthood, although neuron-specific *Nrxn1* cKO mice exhibited reduced body weights (Extended Data Fig. 4c-d). The neuron and the astrocyte *Nrxn1* cKO each decreased the total Nrxn1α protein levels by approximately 50% (Fig. 2c-d). Immunoblotting with antibodies that recognize all neurexins revealed considerable reductions in the total α-neurexin levels in neuron and astrocyte *Nrxn1* cKOs mice, consistent with the high expression levels of Nrxn1 (Fig. 2c, 2e; Extended Data Fig. 4e-f). Deletion of *Nrxn1* in neurons, but not in astrocytes, resulted in a 50% decrease in total β-neurexin levels (Extended Data Fig. 4e-f). Thus, astrocytes and neurons produce similar amounts of Nrxn1α protein in brain, while only neurons express Nrxn1β protein. These data were confirmed with HA-Nrxn1 cKI mice (Extended Data Fig. 4g-i).

Modification of Nrxn1 by glycosylation^23^ and heparan sulfate^26, 57^ has been implicated in controlling ligand binding and synapse formation. Immunoblotting analyses of Nrxn1α in cultures composed of only glia or glia mixed with neurons revealed that glia express Nrxn1α at a much higher molecular weight than neurons (Extended Data Fig. 5a-b). Heparinase treatment of Nrxn1α immunoprecipitated from glia cultures caused a shift of nearly all Nrxn1α protein to a lower molecular weight, whereas the same treatment of Nrxn1α immunoprecipitated from mixed neuron and glia cultures produced only a partial shift (Extended Data Fig. 5a-b). Quantifications of the heparan sulfate ‘stubs’ remaining after heparinase treatment demonstrated that unlike mixed neuron-glia cultures, Nrxn1α derived from glia is almost completely modified by heparan sulfate (Extended Data Fig. 5c).

**Figure 5.**
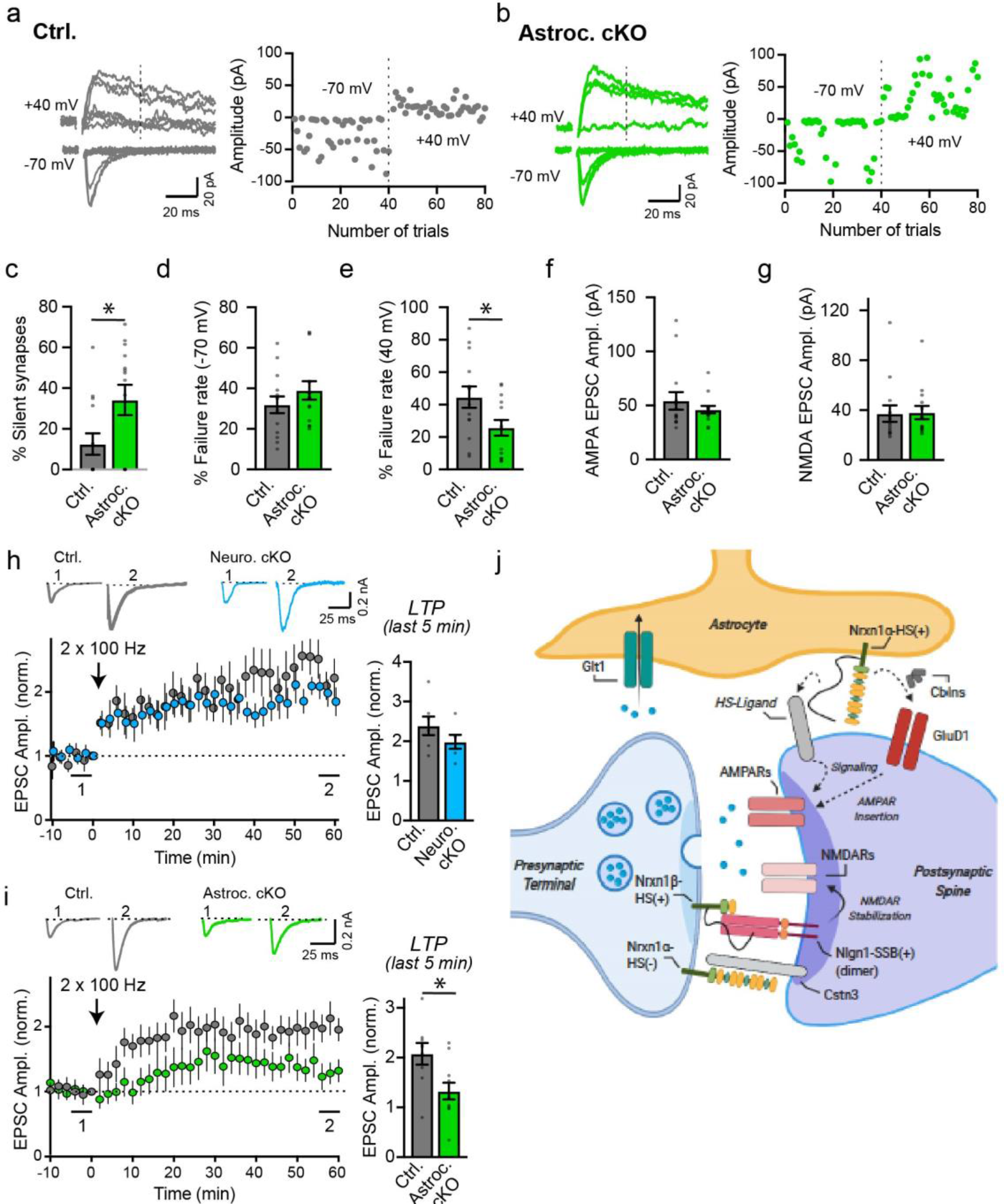
Astrocytes promote synapse unsilencing and long-term potentiation in a Nrxn1-dependent manner. **a-b,** Minimal stimulation recordings were performed on acute hippocampal slices from ctrl (**a**) and Nrxn1 astrocyte cKO mice (**b**). Example traces (left) and scatter plot (right) of evoked EPSCs from CA1 pyramidal neurons at -70 and +40 mV holding potentials. Failed events were assigned an amplitude of 0 pA for visualization. **c**, Astrocyte-specific deletion of Nrxn1 increases the percentage of silent synapses by 3-fold. Cells with EPSCs detected at both holding potentials are indicated on the x-axis (ctrl., 9 cells; astrocyte cKO, 5 cells). **d-e**, Summary graphs of failure rate averages of individuals cells held at -70 (**d**) and +40 mV (**e**) holding potentials. **f-g,** Summary graphs of average AMPA-EPSC (**f**) and NMDA-EPSC (**g**) amplitudes measured at near unitary, minimal stimulation. **h-i,** Neuron-specific deletion of Nrxn1 does not significantly affect the induction of NMDAR-dependent LTP (**h**), whereas the astrocyte-specific deletion of Nrxn1 severely impairs LTP (**i**). Schaffer-collateral LTP was induced by two 100-Hz/1 s stimulus trains separated by a 10 s interval, and recorded from CA1 pyramidal neurons (top, sample traces; bottom left, summary plot of LTP time course; bottom right, LTP amplitude during the last 5 min of recording). **j,** Summary model of presynaptic and astrocytic Nrxn1 signaling within tripartite synapses. All data are means ± SEM. *n =* 14 cells / 7 mice ctrl., 14 cells / 4 mice (**c**-**g**); 7 cells / 8 mice for ctrl., 6 cells / 4 mice for neuron cKO (**h**); 9 cells / 6 mice for ctrl., 11 cells / 7 mice for astrocyte cKO (**i**). Statistical analysis was assessed by two-tailed unpaired t-test with * = p < 0.05.

These results indicate that Nrxn1α is preferentially, if not exclusively, expressed as a heparan sulfate proteoglycan in astrocytes but not in neurons. To validate this conclusion, we analyzed Nrxn1 isoforms in the brains of littermate control mice and of neuron- and astrocyte-specific cKO-T mice (Extended Data Fig. 5d-f). Native brain Nrxn1α migrated on SDS-gels as a heterogeneous set of bands with a diffuse high-molecular weight component. Deletion of Nrxn1 from neurons selectively decreased the intensity of the lower bands, whereas deletion of Nrxn1 from astrocytes nearly eliminated the diffuse high-molecular weight component (Extended Data Fig. 5d). We next treated Nrxn1α immunoprecipitated from the brains of the control and neuron- and astrocyte-specific cKO-T mice with heparinases and analyzed the resulting heparan-sulfate stubs. Deletion of Nrxn1 from neurons did not significantly decrease the levels of heparan sulfate-modified Nrxn1α as analyzed by the heparinase-induced generation of heparan-sulfate stubs, whereas deletion of Nrxn1 from astrocytes almost completely eliminated heparan sulfate-modified Nrxn1α (Fig. 2f-h). Thus, Nrxn1α is primarily heparan-sulfate modified in brain in astrocytes but not neurons. Consistent with this conclusion, Nrxn1 ligands that appear to block the heparan sulfate modification of neurexins (i.e. CA10^58^ and FAM19A1-4^59^) are not expressed at appreciable levels in astrocytes (Extended Data Fig. 5g).

Since astrocytic and neuronal Nrxn1 may converge upon similar postsynaptic structures and share access to ligands, differential alternative splicing of *Nrxn1* mRNAs represents a powerful potential mechanism to direct information flow between different compartments of the tripartite synapse. To determine whether alternative splicing of *Nrxn1* mRNAs differs between neurons and astrocytes, we compared *Nrxn1* mRNA alternative splicing in neurons and astrocytes from brain tissue using mRNA isolated via RiboTag pulldowns (described in Fig. 1a & 2b; Extended Data Fig. 1a). In brain, astrocytic *Nrxn1* mRNA displayed strikingly more restricted patterns of alternative splicing than neuronal *Nrxn1* mRNA (Fig. 2i; Extended Data Fig. 6a-f). In particular, astrocytic *Nrxn1* mRNA nearly always lacked inserts in splice sites #1, #2, and #6, but invariably contained an insert in splice site #4 (Fig. 2i; Extended Data Fig. 6a-f). These observations were confirmed by comparing *Nrxn1* splicing in pure glia vs. mixed neuron-glia cultures (Extended Data Fig. 6g-h).

In heterologous synapse formation assays, neurexin ligands (e.g. neuroligins) expressed on the surface of non-neuronal cells (e.g. COS or HEK293T cells) cluster neurexins on the surface of co-cultured neurons and induce formation of presynaptic specializations on these neurons^60, 61^. We developed a modified version of this assay using pure glia cultures from *Nrxn1* HA-knockin mice to screen ligands for binding to endogenous astrocytic Nrxn1. We found that only a subset of established synaptic Nrxn1 ligands could cluster astrocytic Nrxn1 (Fig. 2j; Extended Data Fig. 6i-j), demonstrating that molecular differences in Nrxn1 isoforms expressed by neurons and astrocytes confer ligand specificity. In particular, we found that astrocytic Nrxn1 could interact with a membrane-anchored variant of neurexophilin-1, neuroligin-1 lacking the splice site B insert (-SSB), and cerebellin-1 complexed with GluD1 (Fig. 2i-j). Collectively, these data provide a molecular basis for segregated, compartment-specific signaling within tripartite synapses by presynaptic neuronal and astrocytic Nrxn1 nanoclusters.

## Astrocytes regulate synapse strength independent of synaptogenesis

The association of molecularly distinct astrocytic and neuronal Nrxn1 nanoclusters with excitatory synapses suggests divergent functions. We next asked whether the neuronal or astrocytic deletion of *Nrxn1* alters synapse numbers, motivated by a large number of studies implicating astrocytes in synaptogenesis^7–15^ and by reports that the heparan sulfate modification of Nrxn1, which we find is relatively specific for astrocytes, is important for synapse formation^26, 57^. Using acute slices from young mice with neuron- and astrocyte-specific *Nrxn1* deletions, we measured spontaneous miniature excitatory postsynaptic currents (mEPSCs) in CA1 pyramidal neurons. Strikingly, we observed a large decrease in mEPSC frequency after astrocyte-specific *Nrxn1* deletions (∼55%), but not after neuron-specific *Nrxn1* deletions (Fig. 3a-b). The decrease in mEPSC frequency induced by deletion of *Nrxn1* from astrocytes would be consistent with a decrease in synapse numbers but could also be due to other causes. Therefore, we measured the synapse density in different layers of the CA1 region by immunohistochemistry for the excitatory presynaptic marker vGluT1 (Fig. 3c-f) and by electron microscopy (Fig. 3g-h). Unexpectedly, we observed no change in synapse density by either method. Furthermore, electron microscopy revealed no significant alteration in the ultrastructure of synapses (Extended Data Fig 7a-f). We also found no significant change in the levels of synaptic proteins (Extended Data Fig 7g-l). Using RNAseq with bulk RNA, we found limited and non-overlapping changes in global gene transcription following deletion of *Nrxn1* in either neurons or astrocytes (Extended Data Fig. 8), consistent with the notion that astrocytic and neuronal Nrxn1 are functionally distinct.

## Neurons and astrocytes promote distinct aspects of functional synapse assembly

The decreased mEPSC frequency observed after astrocyte-specific deletions of *Nrxn1* (Fig. 3b) could be due to a decrease in synaptic transmission rather than synapse numbers. To explore this possibility, we monitored AMPAR- and NMDAR-mediated synaptic responses in acute slices obtained from neuron- and astrocyte-specific *Nrxn1* cKO mice. Using whole-cell patch-clamp recordings, we found that the neuron-specific *Nrxn1* cKO significantly decreased the NMDAR/AMPAR ratio (Fig. 4a), whereas the astrocyte-specific cKO increased the NMDAR/AMPAR ratio (Fig. 4b).

Subsequently, we measured the strength of AMPAR- and NMDAR-mediated synaptic transmission separately, using input-output curves to control for differences in action potential generation. The neuron-specific cKO of *Nrxn1* decreased only the strength of NMDAR-mediated synaptic transmission (Fig. 4c, 4e), while the astrocyte-specific cKO of *Nrxn1* decreased only the strength of AMPAR-mediated synaptic transmission (Fig. 4d, 4f). Impaired synaptic transmission induced by *Nrxn1* deletions in neurons and astrocytes was not due to a decrease in release probability, because the paired-pulse ratio was unchanged (Extended Data Fig 9e-f). Since the release probability appeared to be normal, we determined whether postsynaptic receptor responses were decreased following neuron- and astrocyte-specific deletion of *Nrxn1* by puffing either NMDA or AMPA onto neurons. Importantly, currents elicited by NMDA, but not by AMPA, were reduced following deletion of Nrxn1 in neurons (Fig. 4g & 4i). Conversely, currents elicited by AMPA, but not by NMDA, were decreased by deletion of Nrxn1 in astrocytes (Fig. 4h & 4j). Thus, deletion of the same gene (Nrxn1) in neurons and astrocytes exerts non-overlapping effects on postsynaptic transmission at Schaffer-collateral synapses.

## Astrocytes promote synapse unsilencing and synaptic plasticity in a Nrxn1-dependent manner

The normal mEPSC amplitudes (Fig. 3b) and kinetics (Extended Data Fig. 9c-d) following deletion of *Nrxn1* in astrocytes suggest that the overall AMPAR function is normal, whereas the paired-pulse measurements indicate that the release probability is normal as well (Extended Data Fig. 9f). Why then does the *Nrxn1* deletion in astrocytes selectively decrease AMPAR-mediated evoked EPSCs (Figs. 3-4)? One possibility is that deletion of *Nrxn1* in astrocytes increases the number of silent synapses that lack functional AMPA receptors. The presence of AMPAR-lacking silent synapses in postnatal brain is well-documented, and conversion of silent to ‘speaking’ synapses is thought to be important for circuit development and long-term synaptic plasticity^62–65^. Moreover, the preferential association of astrocytes with larger, morphologically mature synapses^1, 2, 56^ suggests that their function may be particularly important for promoting synapse unsilencing. Using a standard minimal stimulation protocol, we found that the deletion of Nrxn1 in astrocytes produces a large increase (∼3-fold) in the abundance of silent synapses (Fig. 5a-b). At minimal stimulation intensities, which elicit near unitary responses, the average size of AMPAR-mediated currents was normal, thus confirming that AMPARs are normal at non-silenced synapses (Fig. 5f).

A recent study found that chemogenetic manipulation of signaling in astrocytes affects long-term potentiation in the hippocampus^66^. Moreover, hippocampal LTP has been shown to alter the physical association of astrocyte processes with synapses^67, 68^. We thus asked whether astrocytic or neuronal Nrxn1 is required for hippocampal LTP. Consistent with the increase in silent synapses, the astrocyte-specific, but not the neuron-specific, *Nrxn1* deletion impaired long-term potentiation (Fig. 5h-i). Our finding that NMDAR-mediated currents are unaltered following deletion of astrocytic *Nrxn1* suggests that astrocytic *Nrxn1* regulates LTP in a manner distinct from astrocyte-mediated secretion of the NMDAR co-agonist d-Serine^69, 70^. Thus, Nrxn1 performs profoundly different synaptic functions in neuronal and astrocytic compartments of SC-CA1 tripartite synapses.

## DISCUSSION

The molecular mechanisms that enable astrocytes to communicate with neurons within tripartite synapses are poorly understood. In particular, earlier work does not address how and why astrocytes physically enmesh synapses, especially larger morphologically-mature synapses, well after their formation^1, 2, 56^. Though still widely debated, the physical association of astrocytes with mature synapses may serve to augment supply of energy substrates, soak up neurotransmitters to prevent diffusion to neighboring synapses, as well as other functions that are not necessarily mutually exclusive, including bidirectional signaling between astrocytes and neurons. Here, we demonstrate that astrocytes throughout brain express high levels of the presynaptic cell adhesion molecule Nrxn1 (Figs. 1-2; Extended Data Fig. 1-5). Nrxn1 is unlike any protein studied in astrocytes to date owing to its unique capability to communicate with synapses via a rich diversity of postsynaptic ligands. We found that astrocytes target Nrxn1 to both vascular endfeet and to excitatory synapses, where they assemble Nrxn1 into nanoclusters that are distinct from presynaptic Nrxn1 nanoclusters. Underlying the ability of presynaptic and astrocytic Nrxn1 to uniquely communicate with postsynaptic neurons, we describe an unexpected molecular choreography between astrocytes and neurons involving differential expression of major *Nrxn1* isoforms, HS-modification of *Nrxn1* protein, and alternative splicing of *Nrxn1* mRNAs (Fig. 2; Extended Data Figs. 4-6; Summary Cartoon, Fig. 5j). Owing to their molecular differences, neuronal and astrocytic *Nrxn1* display unique ligand-binding profiles (Fig. 2j, Extended Data Fig. 6i-j) and regulate distinct aspects of excitatory synapse function in hippocampal area CA1 (Fig. 3-5; Extended Data Fig. 7-9). In particular, we found that deletion of *Nrxn1* in astrocytes impaired the conversion of silent synapses to functional synapses and inhibited long-term synaptic plasticity (Figs. 4-5). In contrast, neuronal deletion of Nrxn1 impaired NMDAR-mediated synaptic transmission (Fig. 5). Our findings thus show that astrocytes modulate synaptic function via a neurexin-dependent mechanism that differs from the neurexin-dependent mechanism of synapse specification mediated by neurons. To our knowledge, our study is the first to pinpoint a specific role of astrocytes in regulating synapse unsilencing independent of synaptogenesis. Our findings indicate that astrocyte-mediated synapse unsilencing represents an ontogenetic window that is distinct from synapse formation and suggests that astrocytes regulate synapse development at several discrete stages, including synapse formation, synapse unsilencing, synapse pruning, and long-term synaptic plasticity.

## MATERIALS AND METHODS

### Animals

Nrxn1 conditional knockout (cKO) and Nrxn1 HA-knockin mice were generated as described previously^32, 45^. Other mouse lines used in this paper include Baf53b-Cre (Jax, stock# 027826), Aldh1l1-CreERT2 (Jax, stock# 029655), Ai14 tdTomato Cre reporter (Jax, stock# 007914), Ai75 tdTomato reporter (Jax, stock# 025106), and RiboTag mice (Jax, stock# 029977). Breeding strategies were designed to ensure that Cre and reporter alleles, when present, were hemizygous. To minimize the potential for germline recombination, Baf53b-Cre alleles were always carried by female breeders and litters were checked for the presence of germline recombined alleles. Weight and survival of Nrxn1 brain, neuron, and astrocyte cKOs was evaluated at post-natal day 21 (P21). Wild-type CD-1 mice were used as control animals for a subset of immunostaining, RNA localization, and biochemical experiments. All mice were weaned at 20 days of age and housed in groups of 2 to 5 on a 12 hr light/dark cycle with access to food and water *ad libidum*. All procedures conformed to National Institutes of Health Guidelines for the Care and Use of Laboratory Mice and were approved by the Stanford University Administrative Panel on Laboratory Animal Care.

### Tamoxifen Preparation and Injections

Tamoxifen (Sigma, Cat# T5648) stock was prepared by shaking 1 g of tamoxifen in 10 ml of 200-proof ethanol at room temperature (RT) for 15 min, followed by mixing with 90 ml of corn oil (Sigma, C8267) for 1-2 hrs at 37°C. Once fully dissolved, 1-ml aliquots (10 mg/ml tamoxifen) were stored at -20°C until use. Exposure to light was minimized at all times. For experiments using Aldh1l1-CreERT2 mice, mice were injected intraperitoneally on P10 and P11 with 300 μl tamoxifen stock using insulin syringes. To obtain both cKO and littermate controls and controls for off-target effects of tamoxifen, entire litters were injected blind to genotype.

### Plasmids

For clustering assays, Nrxn1 ligands were cloned into mammalian expression vectors with tags that enabled detection of ligand expression on the cell surface (Fig. 2j; Extended Data Fig. 6i). Human LRRTM2 was cloned into the pCMV5 vector and contained a PreProTrypsin signal peptide followed by an N-terminal flag sequence. Mouse neuroligin-1 cDNAs differing at splice site B^71^ were cloned into a lentiviral shuttle vector containing a ubiquitin promoter (i.e. FUW). An N-terminal flag sequence was placed after the native signal peptide. Mouse GluD1 was cloned into FUW by PCR amplification of GluD1 cDNA (ORFeome Collaboration Clones, ID# 100068077) and an N-terminal flag sequence was placed after the native signal peptide. A novel membrane-tethered form of mouse neurexophilin-1 was generated by fusing mature neurexophilin-1 possessing an IgK signal peptide to the stalk region of mouse neurexin-3β in place of the LNS6 domain. A flexible linker sequence and flag sequence was inserted between mature neurexophilin-1 and the stalk region to facilitate proper protein folding. The chimeric fusion construct was cloned into FUW. Mouse cerebellin-1 was cloned into the episomal-type pEB-multi expression vector downstream of a CMV5 enhancer and Chicken actin promotor sequence. A flag sequence was placed in the N-terminus after the signal peptide, and a tandem V5 and polyhistidine sequence was placed at the C-terminus. Calsyntenin-3 was kindly provided by Anne Marie Craig and has been described previously^41^. The calsyntenin-3 construct contains an N-terminal myc sequence, a C-terminal mVenus sequence and deletion of a proteolytic cleavage site to enable better surface expression. Enhanced GFP (EGFP) was used to label transfected HEK293T cells. Empty backbone vectors (e.g. pcDNA3.0, pCMV5, and pEB multi) were used for co-transfection experiments.

### Primary Antibodies

The following antibodies were used at the indicated concentrations (IHC-immunohistochemistry; ICC-immunocytochemistry; IB-immunoblot): purified anti-HA mouse (Biolegend Cat# 901501; 1:500 live surface ICC, 1:500 ICC, 1:500 IHC, 1:1000 IB), purified anti-HA mouse Alexa647-conjugated (Biolegend Cat# 682404; 1:500 IHC), anti-HA rabbit (Cell Signaling Cat#3724; 1:250 ICC), anti-Homer1 rabbit (Millipore Cat# ABN37; 1:1000 IHC) anti-Nrxn1 rabbit (Synaptic Systems Cat# 175103; 1:1000 IB), anti-pan-Nrxn rabbit (Frontier Institute Cat# AF870; 1:500 IB), anti-pan-Nrxn rabbit (homemade, G393; 1:500 IB), anti-pan-Nrxn rabbit (homemade, G394; 1:500 IB), anti-pan-Nrxn rabbit (Millipore Cat# ABN-161-l; 1:1000 IB), anti-laminin-alpha2 rat (Abcam Cat# ab11576; 1:5000 IHC), anti-GFAP mouse (Neuromab Cat# 75-240; 1:1000 IB, 1:1000 ICC, 1:500 IHC), anti-GFAP rabbit (Agilent Cat# 033401-2; 1:1000 ICC, 1:1000 IHC), anti-GFAP chicken (Encorbio Cat# CPCA-GFAP; 1:1000 ICC, 1:1000 IHC), anti-HS-stub 3G10 antibody mouse (Amsbio Cat# 370260-1; 1:1000 IB), anti-vGluT1 (Homemade YZ6089; 1:1000 IHC, 1:1000 IB), anti-MAP2 mouse (Sigma Cat# M1406; 1:1000 IHC), anti-NeuN mouse (Millipore Cat# MAB377; 1:1000 IHC), anti-NeuN rabbit (1:1000 IHC), anti-ß-actin mouse (Sigma Cat#A1978; 1:3000 IB), anti-Synapsins rabbit (Homemade YZ6078; 1:500 ICC, 1:1000 IB), anti-Flag rat (Sigma Cat# SAB4200071; 1:500 surface ICC), anti-Myc rat (Abcam Cat# ab206486; 1:500 surface ICC), anti-VGAT guinea pig (Synaptic Systems Cat# 131005; 1:500 IHC), anti-GluN1 mouse (Synaptic Systems Cat# 114011; 1:1000 IB), anti-GluN2B mouse (Neuromab Cat# 75-101; 1:1000 IB), anti-GluR1 rabbit (Millipore Cat# Ab1504; 1:1000 IB), anti-GluR2 mouse (Neuromab Cat# 75-002; 1:1000 IB), anti-GluR4 (Millipore Cat# Ab1508; 1:1000 IB), anti-PSD-95 mouse (Neuromab Cat# 75-028; 1:1000 IB), anti-GRIP mouse (1:1000 IB), anti-Gad67 mouse (Millipore Cat# mab5406B; 1:1000 IB), anti-SNAP25 rabbit (Homemade P913; 1:500 IB), anti-Nlgn1 mouse (Synaptic Systems Cat# 129111; 1:1000 IB), anti-Nlgn2 rabbit (Synaptic Systems Cat# 129203; 1:1000 IB), anti-Nlgn3 mouse (Synaptic Systems Cat# 129311; 1:2000 IB), anti-GluD1 rabbit (Frontier Institute Cat# GluD1C-Rb-Af1390; 1:2000 IB), anti-CASK mouse (NeuroMab Cat# 75-000; 1:1000 IB). anti-Kir4.1 rabbit (Alomone Cat# APC-035; 1:2000 IB), anti-Glt1 guinea pig (Millipore Cat# AB1783; 1:1000 IB)

### Cell and Tissue Lysis

Primary cultures (neurons and glia or glia only) plated in 24-well plates were isolated with 50–80 µl (per well) of ice-cold complete radioimmunoprecipitation assay (RIPA) lysis buffer containing 150 mM NaCl, 5 mM EDTA, 1% Triton X-100 (tx-100), 0.1% SDS, and 25 mM Tris-HCl, pH 7.6 in addition to 1X cOmplete ULTRA protease inhibitor cocktail (Roche, Cat# 11873580001). Fresh or snap-frozen hippocampal tissue was Dounce homogenized in 0.8-1.2 mL of cRIPA. Lysates were incubated on ice for 20 min and then clarified by centrifugation for 20 min at 13,000 rpm at 4°C. Cleared lysates were stored at −80°C until further processing.

### Heparinase Treatment

Heparinase treatment of immunoprecipitated Nrxn1 was performed as described previously^26^ but with some modifications. Each sample was subjected to two separate immunoprecipitation reactions allowing on-bead incubation of purified Nrxn1 in reaction buffer with or without heparinases. Similar quantities of primary culture lysates (150 μg per reaction; Extended Data Fig. a-c) and brain tissue lysates (500 μg per reaction; Fig. 2f-h) were used. Lysates were incubated O/N with prewashed 1:1 anti-HA agarose beads (Sigma, Cat# A2095) at 4°C. Beads were then washed 1X with ice-cold wash buffer (100 mM NaCl, 20 mM Tris-HCl at pH 7, 1.5 mM CaCl_2_, 1X protease inhibitors) containing 1% tx-100 and twice with wash buffer containing 0.2% tx-100. Beads were then incubated with 100 μl reaction buffer (containing 1X NEB heparinase mix diluted in dH2O, 1X cOmplete ULTRA protease inhibitor cocktail, and 0.2% tx-100) with or without 1 μl each of heparinases-I, -II, and -III (NEB) for 2 hrs at 30°C and 600 RPM. Beads were washed once with wash buffer containing 0.2% tx-100 to get rid of heparinases. Proteins were eluted with 60 ul of 2X Laemmli sample buffer with DTT at 65°C for 10 min.

### Immunoblotting

Protein concentrations were determined with the BCA assay, using a BSA standard curve (Life Technologies Cat#23227). Samples were diluted in Laemmli sample buffer (final concentration: 1X) containing fresh DTT and heated to 95°C for 5 min. To limit protein aggregation caused by heating multi-pass transmembrane proteins (i.e. Glt-1, Kir4.1, and vGluT1), some samples were not heated. Proteins were separated by SDS-PAGE using 4–20% MIDI Criterion TGX precast gels (Bio-Rad). Generally, proteins were transferred onto 0.2 µm pore nitrocellulose membranes for 10 min at 2.5 V using the Trans-blot turbo transfer system (Bio-Rad). For more sensitive detection of neurexin levels, proteins were transferred onto nitrocellulose transfer membrane using a Criterion Blotter (Bio-Rad) with plate electrodes in ice-cold transfer buffer (25.1 mM Tris, 192 mM glycine, 20% methanol) at 80V constant voltage for 1 hr. Membranes were blocked in 5% non-fat milk (Carnation) diluted in PBS or TBS for 1 hr at RT. Membranes were then incubated with primary antibodies diluted in TBST (containing 0.1% Tween-20) overnight at 4°C. Beta-actin was used as a loading control for protein quantifications. Membranes were washed 3 times followed by incubation with secondary antibodies. Combinations of the following IRDye secondary antibodies were used (1:10,000 in TBST with 5% milk): IRDye 800CW donkey anti-mouse (926–32212), IRDye 680LT donkey anti-mouse (926–68022), IRDye 800CW donkey anti-rabbit (926–32213), IRDye 680LT donkey anti-rabbit (926–68023), and IRDye 680LT donkey anti-guinea pig (926-68030) from LI-COR. The Odyssey CLx imaging systems (LI-COR) was used to detect signal and was set to automatic mode for dynamic range detection. Pseudo-colors were applied to the signals, and quantification was performed using Image Studio 5.2. Normalization was performed as described in the figure legends.

### Cell Culture

#### Primary neuron cultures (containing glia)

Hippocampal cultures containing neurons and glia (i.e., mixed) were generated from P0 Nrxn1 cKI mice (Extended Data Fig. 5a-c) or wild-type CD-1 mice (Extended Data Fig. 6a-c). Hippocampi were dissected and mixed regardless of gender. In general, pooling tissue from three to six mice in a given preparation was used to generate cultures. Hippocampi were dissected in ice-cold Hank’s balanced salts solution (HBSS) and kept on ice until digestion with 0.2 μm-filtered 10 U/mL papain (Worthington Biochemical Corporation) in HBSS in a 37°C water bath for 20 min. Hippocampi were washed twice with HBSS and once with plating medium [MEM (Gibco) supplemented with 2 mM L-glutamine (Gibco), 0.4% glucose (Sigma), 2% Gem21 NeuroPlex Serum-Free Supplement (Gemini), and 5% fetal bovine serum (FBS, Atlanta)]. Cells were dissociated by trituration in plating media and plated onto glass coverslips coated with Matrigel Membrane Matrix (Corning) in 24-well plates (1 pup’s hippocampi/12 wells). This day was considered day *in vitro* (DIV) 0. The next morning, on DIV1, 90% of the plating medium was replaced with fresh, pre-warmed growth medium [Neurobasal-A (Gibco) supplemented with 2% Gem21 NeuroPlex Serum-Free Supplement and 2 mM L-glutamine (Sigma)]. At DIV3-4, when glia were ∼70-80% confluent, 50% of the medium was replaced with fresh, pre-warmed plating medium and a final concentration of 2 µM cytosine arabinoside (AraC). At DIV8 and DIV12, 50% of the medium was replaced with fresh, pre-warmed plating medium. Analyses were performed at DIV14-16.

#### Primary glia cultures

Glial cultures were prepared using hippocampi pooled from P1-P2 pups of wild-type CD-1 (Extended Data Fig. 6a-c) or Nrxn1 cKI mice (Figs. 1b, 2j; Extended Data Fig. 5a-c, 6i-j). Tissue was digested with 0.2 μm-filtered 10 U/mL papain (Worthington Biochemical Corporation) in HBSS for 20 min at 37°C, followed by harsh trituration to ensure low numbers of viable neurons^72, 73^. Dissociated hippocampal cells from 2 pups were pooled and plated in T25 flasks. Cells were plated in 90% Dulbecco’s Modified Eagle Medium (DMEM) with 10% FBS and incubated at 37°C and 5% CO_2_. Medium was changed 24 hrs later and supplemented with 0.5% penicillin/streptomycin. Glial cultures were further purified once they reached 90% confluence 7 days later. Cells were rinsed three times, provided with fresh medium, and equilibrated for 2 hrs at 37°C and 5% CO_2_. Flasks were transferred to an orbital shaker (250 RPM) for 15 hrs. After shaking, cells were rinsed three times and provided with fresh media. The cell layer following purification consisted of approximately 95% astrocytes. Re-plating of purified astrocytes was performed by detaching the cells with 0.05% trypsin for 5 min, centrifugation for 5 min at 1200 RPM, resuspending and plating at 30% confluence in 90% DMEM with 10% FBS. Astrocytes were replated on Matrigel-coated coverslips for immunofluorescence experiments or on 12-well culture plates for biochemical analysis. Cultured glia were analyzed once they achieved confluency around DIV14.

#### HEK293T cells

HEK293T cells (ATCC CRL•11268) were grown in complete DMEM (cDMEM), which consisted of DMEM (Gibco), 5% FBS (Sigma), penicillin, and streptomycin. All transfections were performed using lipofectamine 3000 (Invitrogen). For co-culture assays, HEK293T cells were plated on 12-well plates and transfected at ∼90% confluency according to manufacturer’s instructions.

### Single-molecule in situ Hybridization and Immunohistochemistry

Wild-type CD-1 mice were euthanized with isoflurane at P30 followed by transcardial perfusion with ice-cold PBS. Brains were quickly dissected and embedded in Optimal Cutting Temperature (OCT) solution on dry ice. 12-µm thick sections were cut using a Leica cryostat (CM3050-S) and mounted directly onto Superfrost Plus histological slides and stored at −80°C until use. Combined single-molecule FISH and IHC for Nrxn1 mRNA (Advanced Cell Diagnostics, probe cat# 461511-C3) and mouse anti-GFAP antibody (Neuromab, Cat# 75-240) was performed using the multiplex RNAscope platform (Advanced Cell Diagnostics) according to manufacturer instructions for fresh-frozen sections. Briefly, after fixation and soon after *in situ* hybridization steps the slides were washed 2 times for 2 mins each with 1X TBST wash buffer. Slides were then incubated with 10% normal serum (TBS-1% BSA) for 30 mins at RT and proceeded with conventional IHC protocol. Primary and secondary Alexa antibodies were used at 1:500 dilution. Samples were mounted using Prolong Gold antifade mounting medium (ThermoFisher, Cat# P36930).

### Purification of Ribosome-bound mRNA

Ribosome-bound mRNA was purified as described previously^74^ but with modifications. Mice were euthanized using isoflurane and decapitated. Brains were quickly dissected and snap-frozen in liquid nitrogen and transferred to -80°C storage until processing. Frozen brains were partially thawed in fresh homogenization buffer at 10% weight/volume and Dounce homogenized. For Baf53b-Cre / RiboTag mice, hippocampi from a single mouse represented a biological replicate, with 4 mice used in total (2 males, 2 females). For Aldh1l1-CreERT2 / RiboTag mice, hippocampi were pooled from 2 animals for a single biological replicate, and 4 replicates were made in total (2 sets of 2 males and 2 sets of 2 females). Homogenate was clarified by centrifugation and 10% of the supernatant was saved as input. The remaining lysate was incubated with pre-washed anti-HA magnetic beads (Thermo) overnight at 4°C. The beads were washed 3 times with a high-salt buffer followed by elution with RLT lysis buffer with β-ME. Both input and IP samples were subjected to RNA extraction using the QIAGEN RNeasy Micro kit. RNA concentration was determined using a NanoDrop 1000 Spectrophotometer (Thermo) and stored at -80°C until downstream analysis.

### qPCR

For all quantitative RT-PCR experiments, RNA concentration was measured using a NanoDrop and equal quantities of RNA were used to synthesize cDNA with the SuperScript IV First Strand Synthesis Kit (Invitrogen, Cat# 18091050). Equal volumes of cDNA were then mixed with TaqMan Fast Virus 1-Step Master Mix (Thermo) and reactions were performed using the QuantStudio 3 RT-PCR System (Thermo). Transcripts were probed using PrimeTime qPCR Probe Assays (Integrated DNA Technologies), which consist of two primers and one FAM-labeled, ZEN/IBFQ-quenched 5’ nuclease probe. Assays generating C_t_ values >35 were omitted. C_t_ values for technical replicates (duplicate or triplicate) differed by less than 0.5. C_t_ values were averaged for technical replicates. Data were normalized to the arithmetic mean of *ActB* and *Gapdh* using the 2^-ΔΔCt^ method.

For measuring the purity of cell type-specific ribosome-bound mRNA following immunoprecipitation (Figure S2D), the following predesigned assays were used (gene, assay ID): Actb (Mm.PT.51.14022423), Aqp4 (Mm.PT.58.9080805), Sox9 (Mm.PT.58.42739087), PDGF (Mm.PT.56a.5639577), MBP (Mm.PT.58.28532164), P2ry12 (Mm.PT.58.43542033), and Rbfox3 or NeuN (Mm.PT.58.11398454), and vGluT1 or SLC17A7 (Mm.PT.58.12116555). To quantify the mRNA levels of Nrxn isoforms (Fig. 2b), the same mRNAs were probed using the following assays (gene, primer 1, primer 2, probe): Nrxn1αβγ (5’-CGATGTCATCTGTCCCAACA-3’, 5’-GCCATCGGATTTAGCACTGTC-3’, 5’-TGGAGCTGCACATACACCAAGGAA-3’), Nrxn1α (5’-TTCAAGTCCACAGATGCCAG-3’, 5’-CAACACAAATCACTGCGGG-3’, 5’-TGCCAAAAC/ZEN/TGGTCCATGCCAAAG-3’), Nrxn1β (5’-CCTGTCTGCTCGTGTACTG-3’, 5’-TTGCAATCTACAGGTCACCAG-3’, 5’-FAM/AGATATATG/ZEN/TTGTCCCAGCGTGTCCG-3’), Nrxn1γ (5’-GCCAGACAGACATGGATATGAG-3’, 5’-GTCAATGTCCTCATCGTCACT-3’, 5’-ACAGATGAC/ZEN/ATCCTTGTGGCCTCG-3’), Nrxn2α (5’-GTCAGCAACAACTTCATGGG-3’, 5’-AGCCACATCCTCACAACG-3’, 5’-FAM/CTTCATCTT/ZEN/CGGGTCCCCTTCCT-3’), Nrxn2β (5’-CCACCACTTCCACAGCAAG-3’, 5’-CTGGTGTGTGCTGAAGCCTA-3’, 5’-GGACCACAT/ZEN/ACAT CTTCGGG-3’), Nrxn3α (5’-GGGAGAACCTGCGAAAGAG-3’, 5’-ATGAAGCGGAAGGACACATC-3’, 5’-CTGCCGTCA/ZEN/TAGCTCAGGATAGATGC-3’), Nrxn3β (5’-CACCACTCTGTGCCTATTTC-3’, 5’-GGCCAGGTATAGAGGATGA-3’, 5’-TCTATCGCT/ZEN/CCCCTGTTTCC-3’).

For qPCR screen of adhesion molecules (Fig. 1a) and secreted HS-modifying Nrxn1 ligands (Extended Data Fig. 5g) the following qPCR assays were used: GPC6 (Mm.PT.58.11926175), Dag1 (Mm.PT.58.46076316), Cadm1 (Mm.PT.58.11863848), PTPRF (Mm.PT.58.14060589), Nrxn1-all (OG-11^30^), MDGA2 (Mm.PT.58.10917976), NrCAM (Mm.PT.58.11121053), Tenm3 (Mm.PT.58.30978772), Cadm2 (Mm.PT.58.6614872), GPC4 (Mm.PT.58.7577704), Lphn3 (Mm.PT.47.16897577), Flrt1 (Mm.PT.58.45891358), Nrxn2-all (OG19 and OG-119^30^), Nlgn3 (OG-7^30^), BAI3 (Mm.PT.58.21599161), BAI1 (Mm.PT.58.31180263), PTPRS (Mm.PT.58.30580505), Grid1 (Mm.PT.58.32947175), ILRAPL1 (Mm.PT.58.13269289), Clstn1 (Mm.PT.58.6236597), Efnb1 (Mm.PT.58.28819484), NTRK3 (Mm.PT.58.30379995), Lphn2 (Mm.PT.49a.12868555), Nlgn1 (OG-2^30^), DCC (Mm.PT.58.7047849), Lphn1 (Mm.PT.47.6572453), BAI2 (Mm.PT.58.11825400), LRRTM4 (Mm.PT.58.11146838), Lrrc4c (Mm.PT.58.29010435), Nlgn2 (OG-139^30^), Tenm1 (Mm.PT.58.31903272), MDGA1 (Mm.PT.58.32475502), Tenm2 (Mm.PT.58.9415753), Lrrtm2 (Mm.PT.58.6337058.g), Ntng2 (Mm.PT.58.33021394), Tenm4 (Mm.PT.58.17129428), Ntng1 (Mm.PT.58.32073611), Clstn2 (Mm.PT.58.6443231), Cadm3 (Mm.PT.58.11025668), Flrt2 (Mm.PT.58.8519890), Flrt3 (Mm.PT.58.11313356), Lrrtm3 (Mm.PT.58.31131475), Clstn3 (Mm.PT.58.5684120), Nrxn3-all (OG-99^30^), Lrrtm1 (Mm.PT.58.42587284.g), PTPRD (Mm.PT.58.45964964), Nptxr (Mm.PT.58.11296212.g), Car10 (Mm.PT.58.11765793), Car11 (Mm.PT.58.32895602), Fam19a1 (Mm.PT.56a.6079538), Fam19a2 (Mm.PT.58.7298614), and Fam19a4 (Mm.PT.56a.9330679).

For qPCR validation of RNAseq hits (Extended Data Fig. 8), the following assays were used (gene name, assay ID): Syndig1 (Mm.PT.58.11714227), A2m (Mm.PT.58.8228034), C1ql3 (Mm.PT.58.27483950), Reln (Mm.PT.58.10165516), Nptx2 (Mm.PT.58.31290939), Ndnf (Mm.PT.56a.9101884), Gigyf1 (Mm.PT.58.5433296), Gnb2 (Mm.PT.58.5245057.g), Lrch4 (Mm.PT.58.41468474), Slc5a5 (Mm.PT.58.31918684), Hpgd (Mm.PT.56a.9684089), TMEM59I (Mm.PT.58.31528327.g), Vps37a (Mm.PT.58.9759868), Csgalnact1 (Mm.PT.58.32698019), GPR17 (Mm.PT.58.7204101), Mfap3l (Mm.PT.58.43969827), Hapln4 (Mm.PT.58.28564264), LPL (Mm.PT.58.46006099), and Ocel1 (Mm.PT.58.5930413.g). *Nrxn1* mRNA levels were evaluated using the above-mentioned primers, as well as a custom PCR assay recognizing the floxed *Nrxn1* exon. *Gapdh* (4352932E, Applied Biosystems) was used for gene normalization. Validation was performed using the same RNA samples that were used for bulk RNAseq analysis with 4 biological replicates (2 males, 2 females) per genotype.

### Junction-flanking PCR

Alternative splicing of *Nrxn1* mRNA was analyzed using junction-flanking PCR as published previously^75^ but with modifications. The following primers anneal to constitutive exon sequences that flank splice junctions and thus amplify *Nrxn1* mRNA transcripts with or without alternative splice sequences (splice site, forward primer, reverse primer): SS1 (5’-GGCAAGGACTGCAGCCAA-3’, 5’-ATCGCTGCTGCTTTGAATGG-3’), SS2 (5’- TGGGATCAGGGGCCTTTGAAGCA-3’, 5’-GAAGGTCGGCTGTGCTGGGG-3’), SS3 (5’-GTGTGAAACTCACGGTCAATCTA-3’, 5’-GTGCCATTCATTATCATTGAGGTTATAG-3’), SS4 (5’-CTGGCCAGTTATCGAACGCT-3’, 5’-GCGATGTTGGCATCGTTCTC-3’), SS5 (5’-TTGACCCCTGTGAGCCGAGC-3’, 5’-GGCTGCTGCGACAATCCCCA-3’), and SS6 (5’-AGGCTTTCAAGGTTGCCTGGCA-3’, 5’-CCCATTGCTGCAAGCAAACGCC-3’).

Splice-junction PCR was performed on cDNA synthesized from equal amounts of immunoprecipitated mRNA and total input RNA (Fig. 2i, Extended Data Fig. 6a-h). Splice-junction PCR was also performed on RNA purified from two-week old mixed hippocampal cultures (neurons and glia) and pure glia cultures (Extended Data Fig. 6g-h). Owing to relative differences in molecular weight of PCR products, running conditions were optimized to allow ideal resolution and band separation for quantification. Samples separated on 20% PAGE gels were stained using PAGE GelRed Nucleic Acid Gel Stain (Biotium). Homemade gels using standard agarose or MetaPhor Agarose were stained using GelRed. Stained gels were imaged at sub-saturation using the ChemiDoc Gel Imaging System (Bio-Rad). Quantification was performed using Image Lab (Bio-Rad) or ImageStudioLite (LI-COR). Intensity values were normalized to the size of DNA products to negate intensity differences related to increased dye incorporation with increased DNA length.

### Heterologous Synapse Formation Assay and Astrocyte Co-Culture Assay

The heterologous synapse formation assay (Fig. 2j; Extended Data Fig. 6i-j) was performed as described previously^76^ but with modifications. HEK293T cells were plated in 12-well plates and transfected with Nrxn1 ligands and GFP plasmids using Lipofectamine 3000. At 16-24 hrs post-transfection, cells were lifted using DPBS containing 2 mM EDTA, spun down at 1200 RPM for 5 min and resuspended in neuronal growth media. In order to not overwhelm primary cultures, 100 μl of a 1:60-1:80 dilution of transfected HEK293T cells (per condition) was added per well of primary culture. Fresh cytosine arabinoside (AraC) was added to limit cell proliferation. For the astrocyte co-culture assay, mixed glia at 70-90% confluency (usually around DIV14) were co-cultured with HEK293T cells for 48 hrs and then cells were live-labeled for surface HA-Nrxn1 and fixed. Staining procedures are described in more detailed below.

### Immunocytochemistry

For live surface-labeling experiments (Figure 1b, 2j; Extended Data Fig. 6i-j), cultures were washed once with ice-cold Dulbecco’s PBS containing 1mM MgCl_2_ and 1 mM CaCl_2_ (PBS-MC). Cultures were then incubated on ice with mouse anti-HA (BioLegend) diluted 1:500 in PBS-MC for 30 min. Cultures were gently washed 3 times with cold PBS-MC and fixed for 20 min at RT with 4% (wt/vol) PFA. Following fixation, cultures were washed 2-3 times with DPBS. For surface-labeling experiments, cultures were blocked for 1 h at RT with antibody dilution buffer (ADB) without tx-100(-), which contains 5% normal goat serum diluted in DPBS. Cells were then labeled with Alexa Fluor-conjugated secondary antibodies (1:1000; Invitrogen) diluted in ADB(-) for 1 h at RT. For clustering assay experiments, cells were washed 3 times with DPBS and then briefly post-fixed for 10 min at RT with 4% PFA. They were then incubated with anti-Myc/Flag antibodies to label surface Nrxn1 ligands overnight at 4C followed by 3 washes, incubation with anti-Rat Alexa secondary antibodies, and 3 final washes.

For all immunocytochemistry experiments, cells were washed and then permeabilized and blocked for 1 h with ADB with tx-100(+) which contains 0.3% tx-100 and 5% normal goat serum diluted in DPBS. Non-surface primary antibodies were diluted in ADB(+) and cells were incubated in the cold-room overnight or for 2 hrs at RT. Cultures were washed three times with DPBS and then incubated with Alexa Fluor-conjugated secondary antibodies (1:1000; Invitrogen) diluted in ADB(+) for 1 h at RT. After three additional washes, coverslips were inverted onto glass microscope slides with Fluoromount-G mounting media (Southern Biotech).

### Immunohistochemistry

Mice were anesthetized with isoflurane and then transcardially perfused (∼1 ml/min) for 1 min with 0.1M DPBS (RT) followed by 7 min with 4% PFA (Electron Microscopy Services). For HA-Nrxn1 labeling (Fig. 1c-l; Extended Data Fig. 2a-c, 3), brains were post-fixed for 13-15 minutes at RT with 4% PFA. To ensure adequate fixation, 2 mm thick sagittal and coronal sections were prepared using young brain slicers (Zivic Instruments). Two-hour post-fixation was used to label sparsely deleted Nrxn1 (Extended Fig. 2a-b). For all other stains (Fig. 3c-d) brains were post-fixed in 4% PFA overnight at 4°C. Brains were washed 3 times with DPBS and cryoprotected by a 24-48 hr incubation in 30% sucrose w/v in DPBS. Brains were embedded in OCT Compound (Sakura), sectioned on the sagittal plane at 30-µm using a cryostat, and stored as floating sections in DPBS. For staining, free-floating sections were incubated with blocking buffer (containing 5% NGS and 0.3% tx-100 in DPBS) for 1 hr at RT. Sections were then incubated with primary antibodies diluted in blocking buffer overnight at 4°C on a rocker. After 3 washes, sections were incubated with Alexa dye secondary antibodies diluted in blocking buffer for 1-2 hrs at RT. Sections were washed 3-4 times and then mounted on charged glass slides. After drying, sections were dipped in water and allowed to dry again. For confocal imaging, per slide, 4 droplets of Fluormount-G with or without DAPI was added, slides were coverslipped, and nail polish was used to secure the coverslip until mounting medium hardened.

### Confocal Microscopy

All confocal images were acquired at RT using an inverted Nikon A1RSi confocal microscope equipped with a 20x, 60x, or 100x objective (Apo, NA 1.4) and operated by NIS-Elements AR acquisition software. In general, high magnification images were taken at 1,024 × 1,024 pixels with a z-stack distance of 0.3 µm. Low magnification images were taken at 1,024 x 1,024 pixels with Nyquist recommended step size. Line averaging (2X) was used for most images. Images were acquired sequentially in order to avoid bleed-through between channels. Imaging parameters (i.e., laser power, photomultiplier gain, offset, pinhole size, scan speed, etc.) were optimized to prevent pixel saturation and kept constant for all conditions within the same experiment. Images were analyzed using NIS-Elements Advanced Research software (Nikon). 3D Deconvolution was performed using recommended parameters for 100X images (Fig. 1f-h; Extended Data Fig. 2b, 3a). For quantitative analysis, imaging of brain tissue involved imaging 2 regions of interest (Fig. 3c-d) or each synaptic subfield outlined (Extended Fig. 2a-c) from at least 5 sections per animal. For all IHC stains, following background subtraction, intensities were averaged per animal in a given brain region or subregion and a minimum of 3 animals per genotype were analyzed. All IHC data was collected and analyzed blindly.

### Slide Scanning Microscopy

*Nrxn1* neuron cKO-T brain slices (coronal and sagittal) were labeled for Nrxn1 (Fig. 1c-d) using purified HA-647 (mouse, 1:500, Biolegend, Cat# 682404) as described above. Slide-mounted sections were scanned at 20X using the Olympus VS200 Slide Scanner. The brightness and contrast of images were adjusted based on non-transgenic controls and exported using OlyVIA image viewer software (Olympus).

### Direct Stochastic Optical Reconstruction Microscopy (dSTORM)

dSTORM images were acquired using a Vutara SR 352 (Bruker Nanosurfaces, Inc., Madison, WI) commercial microscope based on single molecule localization biplane technolog^77, 78^. Thirty micrometer thick hippocampal sections were prepared as described and labeled with Homer1 (rabbit, 1:1000, Millipore, Cat#: ABN37) and purified HA-647 (mouse, 1:500, Biolegend, Cat# 682404) primary antibodies and secondary antibodies conjugated to CF568 (1:3000, Biotium). A few additional modifications were included to improve the specificity of labeling for dSTORM. Prior to labeling sections were treated for 15 minutes with 0.1M glycine. Moreover, all solutions were filtered prior to use using 0.2 μm syringe filters and wash buffers included 0.05% tx-100. The slices were mounted on a coverslip coated with poly-L-Lysine. After drying, sections were briefly re-hydrated and post-fixed using 2% PFA for 10-15 minutes followed by three washes with DBS. Sections were stored at 4C in DPBS shielded from light, until imaging. Coverslips were placed in dSTORM buffer containing (in mM) 50 Tris-HCl at pH 8.0, 10 NaCl, 20 MEA, 1% β-mercaptoethanol, 10% glucose, 150 AU glucose oxidase type VII (Sigma Cat#: G2133), and 1500 AU catalase (Sigma Cat#: C40). Labeled proteins were imaged with 647 and 561 nm excitation power of 40 kW/cm2. Images were recorded using a 60×/1.2 NA Olympus water immersion objective and Hamamatsu Flash4 sCMOS camera with gain set at 50 and frame rate at 50 Hz. Data was analyzed by Vutara SRX software (version 6.04). At least two ROI’s were acquired from a minimum of two sections per animal. Single molecules were identified in each frame by their brightness after removing the background. HA-Nrxn1 was imaged for 20,000 frames with the first 3000 frames excluded from analysis. Homer1 was imaged for 4000 frames with the first 1000 frames excluded from analysis. Identified molecules were localized in three dimensions by fitting the raw data in a 12 × 12-pixel region of interest centered around each particle in each plane with a 3D model function that was obtained from recorded datasets of fluorescent beads. Fit results were filtered by a density based denoising algorithm to remove isolated particles and rendered as 50 nm points. The remaining localizations were classified into clusters by density-based spatial clustering of applications with noise (DBSCAN), a minimum of 10 localizations were connected around a 100 nm search radius for Nrxn1 and 50 localizations around a 300 nm search radius for Homer1. A standard 100 nm hull was used for both. Data was filtered using STORM RLA (settings: 20 px/um, no limit, 200 monte carlo, 1000 nm radius, 100 bins). The experimentally achieved image resolution of 40 nm laterally (x, y) and 70 nm axially (z) was determined by Fourier ring correlation.

### Transmission Electron Microscopy

Three pairs of P26 Nrxn1 astrocyte cKO mice and littermate controls (2 males and 1 female per genotype) were perfused with PBS followed by 4% paraformaldehyde w/v in PBS. The brains were dissected out and post-fixed in 2.5% glutaraldehyde and 2% paraformaldehyde in 0.1M sodium cacodylate buffer (pH 7.4) overnight at 4°C. The next day, 200-μm vibratome sections were collected in 0.1M cold cacodylate buffer and were re-fixed in 1% glutaraldehyde in 0.1M cacodylate buffer before being shipped to Yale CCMI EM facility. Hippocampal regions were dissected out and further post-fixed in 1% OsO_4_, 0.8% potassium ferricyanide in 0.1 M cacodylate buffer at RT for 1 hour. Specimens were then *en bloc* stained with 2% aqueous uranyl acetate for 45 min, dehydrated in a graded series of ethanol to 100%, substituted with propylene oxide and embedded in EMbed 812 resin. Sample blocks were polymerized in an oven at 60°C overnight. Thin sections (60 nm) were cut by a Leica ultramicrotome (UC7) and post-stained with 2% uranyl acetate and lead citrate. Sections were examined with a FEI Tecnai transmission electron microscope at 80 kV of accelerating voltage, digital images were recorded with an Olympus Morada CCD camera and iTEM imaging software. Approximately 30 electron micrographs were collected at random and analyzed for each animal in the stratum radiatum of CA1. Image analysis was performed using ImageJ while blind to genotype. Asymmetric excitatory synapses were identified based on the presence of a post-synaptic density and at least two presynaptic vesicles at the active zone. In addition, several morphological features of excitatory synapses were measured including postsynaptic area, PSD thickness, PSD length, number of synaptic vesicles per synaptic terminal, number of docked vesicles per synaptic junction, and width of the synaptic cleft. For statistical comparisons, density measurements were averaged per image and then averaged per animal. Synapse morphology measurements were only averaged per animal.

### RNA Sequencing

Bulk hippocampal transcriptomic analysis was performed on Nrxn1 astrocyte and neuron cKO mice and littermate controls at P26-P30. To minimize batch effects and technical variability, 2 separate rounds of sequencing were performed on samples with 2 mice per genotype per round. In total, each genotype included 2 males and 2 females. Hippocampi were dissected and immediately mechanically homogenized in RLT Plus with β-ME followed by centrifugation through a QIAshredder. RNA was extracted from the flow-through using the RNeasy Plus Mini Kit (QIAGEN). RNA was aliquoted into at least three tubes to minimize freeze-thaw cycles while allowing for QC analysis using a Bioanalyzer, RNA sequencing, and post-sequencing qPCR validation of differentially expressed genes. Library preparation, RNA and library quality measurements, and one-end sequencing was performed at a depth of 30M reads by BGI in Hong Kong, China.

#### Analysis

The R package DESeq2 was used to analyze differentially expressed genes using a negative binomial distribution, as described previously^79^. The DESeq data set was prepared based on the matrix of raw, un-normalized counts. The DESeq2 model internally corrected for library size, and the mean expected counts of each gene within each group were adjusted by normalization factors. As two separate rounds of sequencing were performed to account for batch variation, batch effects were controlled for in the design formula associated with the DESeq2 data set by using “batch” and “condition” variables to model the samples. Independent filtering of the mean normalized counts was performed using the “results” function, where a False Discovery Rate (FDR) threshold for adjusted p-values was set to alpha < 0.1. Genes of interest for subsequent qPCR validation were determined based on expression changes greater or less than 15% and p-values of less than 0.001, as well as adjusted p-values generally less than 0.05. Several genes with adjusted p-values between 0.2 and 0.05 were also tested. Volcano plots were generated using the VolcaNoseR package (https://huygens.science.uva.nl/VolcaNoseR/).

### Electrophysiology

Acute transverse brain slices (300 μm) containing the dorsal hippocampus were prepared from P22-26 Nrxn1 brain, astrocyte, and neuron cKO mice and their WT littermates. Mice were anesthetized with isoflurane and decapitated. The brain was rapidly removed and placed into ice cold cutting solution containing the following (in mM): 205 sucrose, 2.5 KCl, 1.25 NaH_2_PO_4_, 25 NaHCO_3_, 25 glucose, 0.4 ascorbic acid, 1 CaCl_2_, 2 MgCl_2_ and 3 sodium pyruvate in double-distilled water (ddH2O), pH 7.4 with NaOH, osmolality 300-305 mOsm saturated with 95% O2/5% CO_2_. Slices recovered at 31 °C in a 50:50 mixture composed of cutting saline and artificial cerebrospinal fluid (aCSF) containing the following (in mM): 123 NaCl, 2.5 KCl, 1.25 NaH_2_PO_4_, 25 NaHCO_3_, 25 glucose, 2 CaCl_2_ and 1 MgCl_2_ in ddH2O and saturated with 95% O2/5% CO_2_, pH 7.4 with NaOH, osmolarity 300-305 mOsm for 30 min and then placed at RT in oxygenated aCSF alone for 1 hr. Whole-cell voltage clamp recordings were performed on CA1 pyramidal neurons. Brain slices were maintained at ∼30°C via a dual-T344 temperature controller (Warner Instruments). Brain slices were continuously perfused with normal oxygenated aCSF (at about 1 ml/min perfusion rate) throughout recordings. Electrical signals were recorded at 25 kHz with a two channel Axoclamp 700B amplifier (Axon Instruments), digitalized with a Digidata 1440 digitizer (Molecular devices) that was in turn controlled by Clampex 10.7 (Molecular Devices). Recording pipettes were pulled from thin-walled borosilicate glass pipettes to resistances of 3-5 MΩ. mEPSCs and EPSCs were recorded with an internal solution containing (in mM): 117 Cs-methanesulfonate, 15 CsCl, 8 NaCl, 10 TEA-Cl, 0.2 EGTA, 4 Na2-ATP, 0.3 Na2-GTP, 10 HEPES, and 2 QX-314 pH pH 7.3 with CsOH (∼300 mOsm). mEPSCs were recorded in 1 μM tetrodotoxin (TTX) and 100 μM picrotoxin. Miniature events were handpicked and analyzed in Clampfit 10 (Molecular Devices) using template matching and a threshold of 5 pA.

Evoked synaptic currents were elicited with a bipolar stimulating electrode (A-M Systems, Carlsborg, WA), controlled by a Model 2100 Isolated Pulse Stimulator (A-M Systems, Inc.), and synchronized with the Clampfit 10 data acquisition software (Molecular Devices). AMPA-receptor-mediated EPSCs were recorded at holding potentials of −70 mV, whereas NMDA-receptor-mediated EPSCs were recorded at +40 mV and quantified at 50 ms after the stimulus artifact. Measurements of the AMPAR/NMDAR ratio, AMPAR and NMDAR input-output curves were performed in 100 μM picrotoxin. Paired-pulse ratios were monitored with interstimulus intervals of 20-200 ms. LTP measurements in CA1 pyramidal neurons were performed using whole-cell patch-clamp recordings with the same extracellular and internal solutions, except the extracellular solution included picrotoxin (100 μM). Schaffer-collateral axons were stimulated extracellularly, and LTP was induced by two applications of 100 Hz stimuli separated by 10 s under voltage-clamp mode (holding potential = 0 mV). Pre-LTP (averaging last 5 mins as baseline) and post-LTP (averaging the last 5 mins) were recorded at 0.1 Hz. For all recordings and analyses, the experimenter was blind to genotype. Electrophysiological data were analyzed using Clampfit 10.4 (Molecular Devices).

Puffing applications of AMPA (50 µM, R-S AMPA hydrobromide, Tocris Bioscience, in the presence of picrotoxin, D-APV and TTX) and NMDA (50 µM, Tocris Bioscience, in the presence of picrotoxin, CNQX and TTX) were performed with 10 psi for 5-200 ms by using Picospritzer III (Parker Instrumentation). The total charge was calculated within 10 s, 20 s, 40 s from puff application. The slope was calculated from the data obtained between the pre-puffing baseline to 100 ms for AMPA and to 50 ms for NMDA.

For minimal stimulation experiments, 250 μm coronal brain slices were prepared. Cutting solution contained the following (in mM): 228 sucrose, 2.5 KCl, 1 NaH_2_PO_4_, 26 NaHCO_3_, 11 glucose, 0.5 CaCl_2_, 3 MgCl_2_ in double-distilled water (ddH2O), pH 7.4 with NaOH, osmolality 300 mOsm. aCSF contained the following (in mM): 119 NaCl, 2.5 KCl, 1 NaH_2_PO_4_, 26 NaHCO_3_, 11 glucose, 2 CaCl_2_ and 1 MgCl_2_ pH 7.4 with NaOH, osmolarity 290-295 mOsm. Recordings were performed in aCSF containing 100 μM picrotoxin with an internal solution containing (in mM): 135 Cs-methanesulfonate, 8 NaCl, 0.3 EGTA, 2Mg2-ATP, 0.3 Na2-GTP, 10 HEPES, 0.1 spermine, 7 phosphocreatine and 5 QX-314 pH pH 7.3 with CsOH (∼300 mOsm). 40-60 AMPA-receptor-mediated EPSCs were recorded at holding potentials of −70 mV. The stimulation intensity was adjusted to obtain a failure rate close to 50%. 40-60 NMDA-receptor-mediated EPSCs were recorded at the same stimulation intensity at holding potentials of +40 mV and quantified 30 ms after the stimulus artifact. The fraction of silent synapses was calculated using the following calculation: 1-Ln(Frequency-80mV)/Ln(Frequency+40mV)

### Quantification and Statistical Analysis

Quantifications have been described in the respective materials and methods sections, and statistical details are provided in the figure legends. Statistical significance between various conditions was assessed by determining p-values (95% confidence interval). For animal survival analysis and quantitative immunoblotting experiments, statistical analyses were performed using GraphPad Prism 6 software. Blot intensity quantification was performed using Image Studio Lite. For most staining experiments, the “n” represents the average per animal or average per culture. In contrast, for electrophysiology measurements, the “n” represents the total number of cells patched. For biochemical and behavioral experiments, the “n” generally represents number of animals, independent cultures or pooled samples. Most intergroup comparisons were done by two-tailed Student’s t test or the Mann Whitney Test. For multiple comparisons, data were analyzed with one- or two-way ANOVA followed by a post-hoc test (e.g. Tukey’s). Levels of significance were set as * p < 0.05; ** p < 0.01; *** p < 0.001; **** p < 0.0001. All graphs depict means ± SEM.

## ACKNOWLEDGEMENTS

We would like to thank Ben Barres, Shane Liddelow and other members of the Barres lab family for discussions, reagents and technical assistance. We thank Ann Marie Craig for the Calsyntenin-3 construct and Anna Kalaj for feedback and discussions. **Funding:** This study was supported by grants from NIMH (MH052804 and MH104172 to T.C.S.; by F32-MH105040, K01-MH123788, and BBRF to J.H.T.).

## AUTHOR CONTRIBUTIONS

J.T. performed the genetic, biochemical, and imaging experiments, Z.D. most electrophysiology experiments, A.S. performed the silent synapse electrophysiology, K.R. contributed to Ribotag experiments, S.E.P. contributed to culture and qPCR experiments, A.N. contributed to RNAseq experiments, X.L. performed the electron microscopy, and K.L. performed the *in situ* hybridizations and contributed to the splicing experiments. J.T. and T.C.S. conceived and planned the project, analyzed the data, and wrote the paper with input from all authors.

## EXTENDED DATA FIGURES and LEGENDS

**Extended Data Figure 1.**
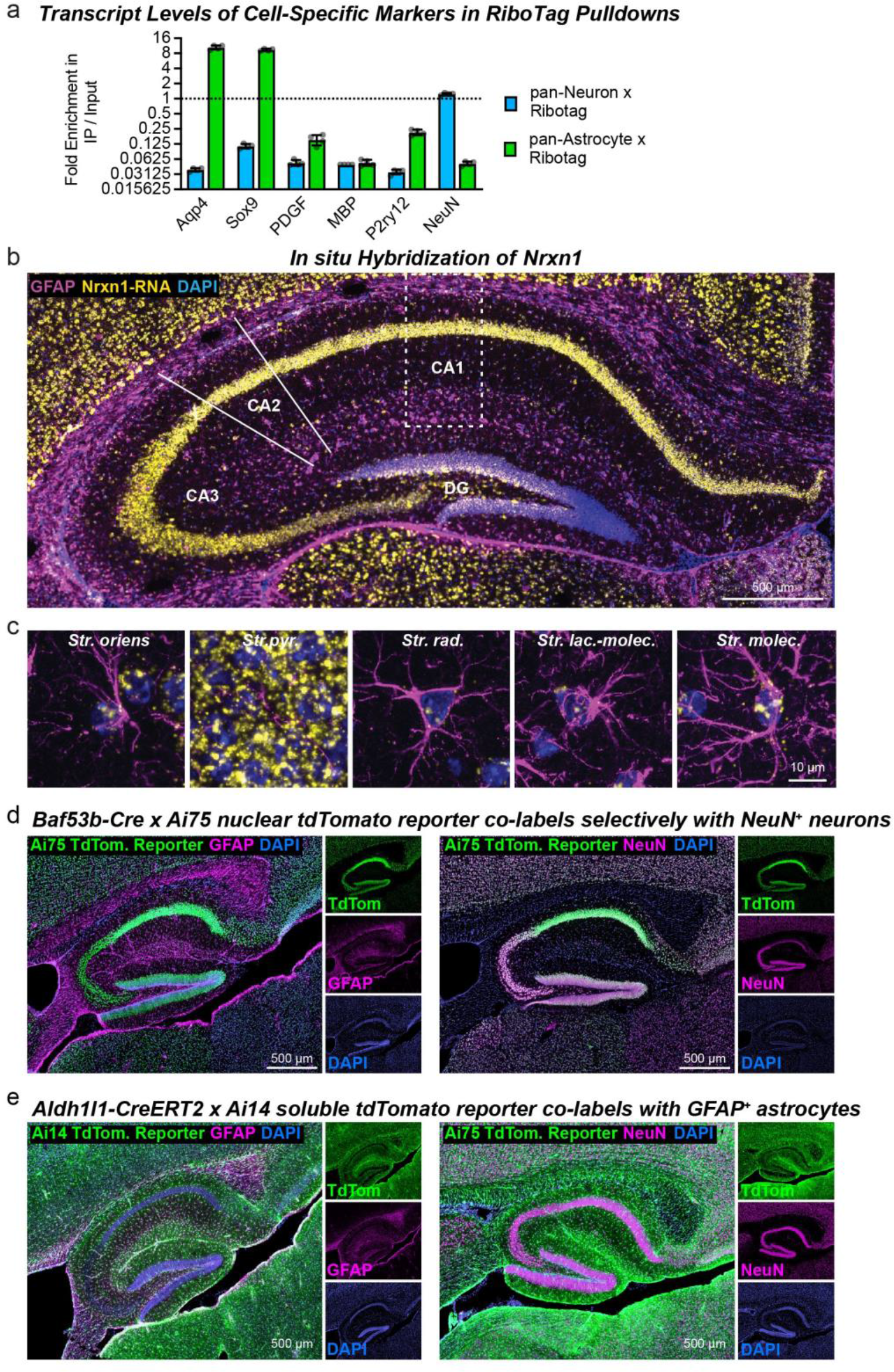
Confirmation of RiboTag pulldown purity by quantitative RT-PCR (a), *In situ* hybridization for *Nrxn1* in the hippocampus, and validation of neuron- and astrocyte-specific Cre driver lines by use of Cre-dependent reporter mouse lines (d-e; data complement those shown in Fig. 1). **a,** Quantifications of cell-specific markers with expected enrichment of astrocyte marker genes (i.e. Aqp4 and Sox9) and a neuron marker gene (i.e. RBFOX3) in astrocyte- or neuron-specific ribosome-bound mRNAs, respectively. The expected de-enrichment in a variety of non-targeted cell-specific markers is seen. Ribosome-bound mRNAs were isolated by immunoprecipitation from P26-P30 hippocampal homogenates of RiboTag mice crossed with neuron- or astrocyte-specific Cre-driver lines. The mRNA abundance (normalized to ActB) of cell markers was compared between immunoprecipitated mRNA (IP) or total mRNA (input) following quantitative RT-PCR. **b-c,** Hippocampal section from young adult mouse (P30) were labeled by single-molecule *in situ* hybridization for *Nrxn1* (yellow), by immunocytochemistry for GFAP (purple), and by DAPI staining for nuclei (blue). Representative images show the entire hippocampal formation (**b**) and expanded images (**c**) of layer specific astrocytes in the CA1 region (*Str. oriens*, *Str. pyramidale*, *Str. radiatum*, *Str. lacunosum moleculare*) and dentate gyrus (*Str. molecular*). **d,** Validation of neuron-specific Cre-recombinase expression by Baf53b-Cre mice^1^ that were crossed with Ai75 TdTomato reporter mice, which express nuclear-targeted tdTomato (shown in a green false color) following Cre recombination. Brain sections from P26 mice were stained with DAPI (blue) to label nuclei, and either with antibodies for GFAP to label astrocytes (left panels, magenta) or antibodies for NeuN to label neurons (right panels, magenta). NeuN(+) cells are tdTomato-positive while astrocytes lack reporter expression. **e,** Validation of astrocyte-specific Cre-recombinase expression by tamoxifen-inducible Aldh1l1-CreERT2 mice^2^ that were crossed to the Ai14 TdTomato Cre reporter line, which expresses cytoplasmic tdTomato (labelled in a green false color). Mice were injected with tamoxifen on P10-P11 and analyzed at P26. Staining was performed as in **d,** revealing selective Cre expression by astrocytes and not neurons. Data are means ± SEM. *n =* 4 mice for neuron mRNAs and 8 mice with 2 mice pooled per pull-down for astrocyte mRNAs (**a**).

**Extended Data Figure 2.**
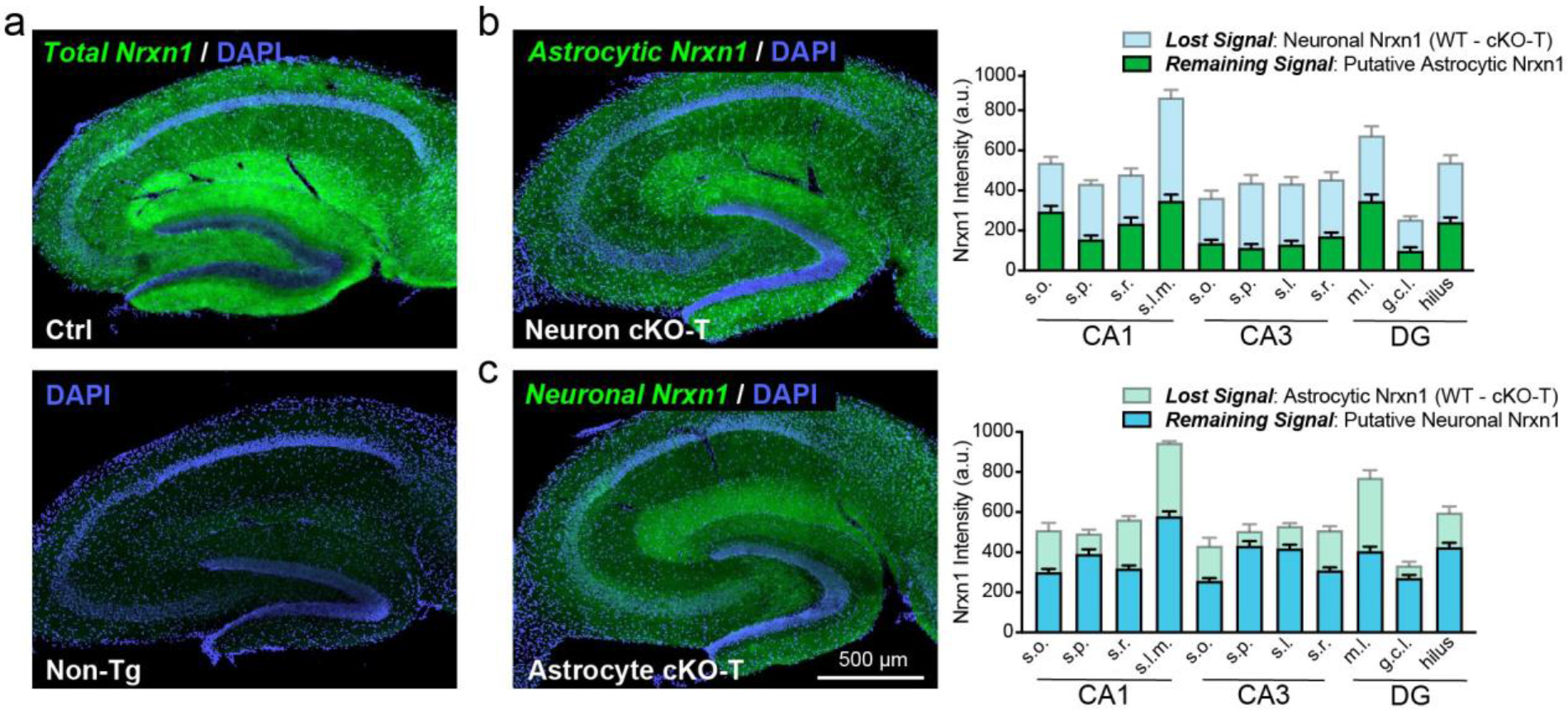
Astrocytes Contribute Significantly to Nrxn1 Levels in Hippocampal Neuropil (data complement those shown in Figure 1). **a**, Nrxn1 (green) was visualized using a dye-conjugated HA antibody in lightly fixed hippocampal slices (i.e. 15 minute post-fixation). Nuclei (blue) were stained using DAPI. Ctrl mice (top) have abundant and layer-specific labeling in the hippocampus, which is entirely absent in sections from non-transgenic mice (bottom). **b-c**, Representative images (left) and summary graphs (right) of HA-Nrxn1 levels detected in sections from Nrxn1 neuron cKO-T (**b**) and astrocyte cKO-T mice (**c**). Background was subtracted based on fluorescence levels detected in labelled non-transgenic sections. Nrxn1 levels were then quantified in major synaptic subfields of CA1 (*Stratum oriens* or *s.o*., *Stratum pyramidale* or *s.p*., *Stratum radiatum* or *s.r*., and *Stratum lacunosum moleculare* or *s.l.m*.), CA3 (*s.o*., *s.p*., *Stratum lucidum* or *s.l*., and *s.r*.), and the dentate gyrus (DG; *molecular layer* or *m.l*., granule cell layer or g.c.l., and the hilus). Remaining Nrxn1 levels in neuron cKO-T (**b**) and astrocyte cKO-T mice (**c**) are considered to be putative astrocyte and neuron Nrxn1 signal, respectively. The loss of signal corresponding to the cKO-T mice is calculated by subtracting cKO-T levels from WT levels. Data are means ± SEM. *n =* 8 sections / 3 WT mice, 11 sections / 4 neuron cKO mice (**b**); 9 sections / 3 WT mice, 18 sections / 6 astrocyte cKO mice (**c**).

**Extended Data Figure 3.**
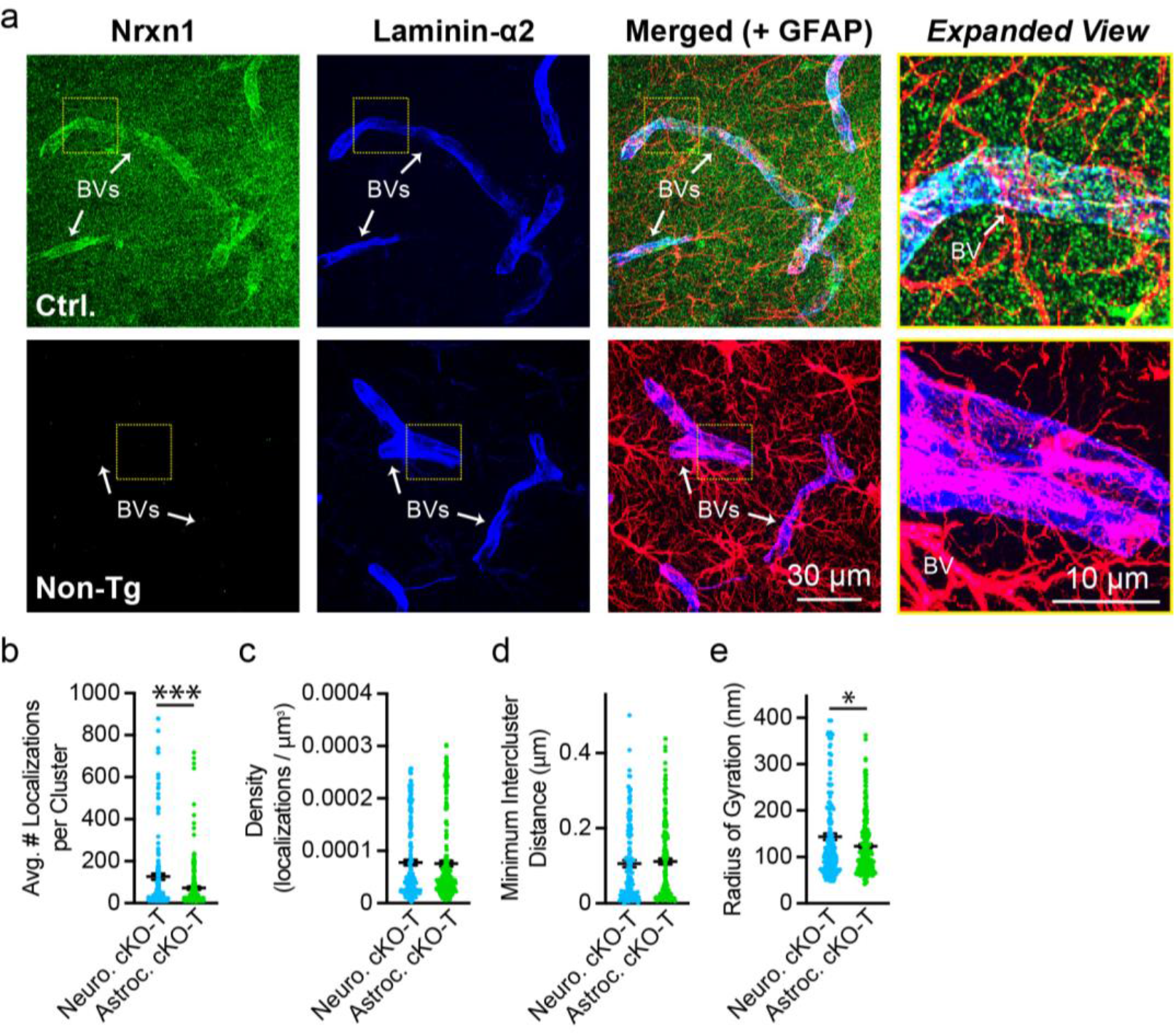
Control stains for Nrxn1 labeling of blood vessels (BVs) (a) and comparison of Nrnx1 nanocluster properties in Nrxn1 neuron and astrocyte cKO-T mice (b-e; data complement those shown in figure 1). **a**, Representative images of Nrxn1 labeling in sections from HA-Nrxn1 cKI (i.e. Ctrl) and non-transgenic mice. Brains were lightly fixed (i.e. 15 minutes post-fixation) and stained for HA-Nrxn1 (green) with dye-conjugated primary antibody, GFAP (astrocytes, magenta), and laminin-alpha2 (blood vessels, blue). Yellow box, enlarged on right. **b-e**, Summary graphs comparing the properties of putative astrocytic Nrxn1 nanoclusters (found in neuron cKO-T mice) and presynaptic Nrxn1 nanoclusters (found in astrocyte cKO-T mice). Properties measured include the average number of localizations per Nrxn1 nanocluster (**b**), the density of localizations per Nrxn1 nanocluster (**c**), the minimum intercluster distance between Nrxn1 and Homer1 clusters (**d**), and Nrxn1 nanocluster size measured as the radius of gyration (**e**). Numerical data are means ± SEM. *n* = 190 synapses / 3 neuron cKO-T mice, 227 synapses / 3 astrocyte cKO-T mice (**b-c, e**) and 130 synapses / 3 neuron cKO-T mice, 190 synapses / 3 astrocyte cKO-T mice (**d**). Statistical significance was determined by two-tailed unpaired t-test to controls with * = p < 0.05; *** = p < 0.001.

**Extended Data Figure 4.**
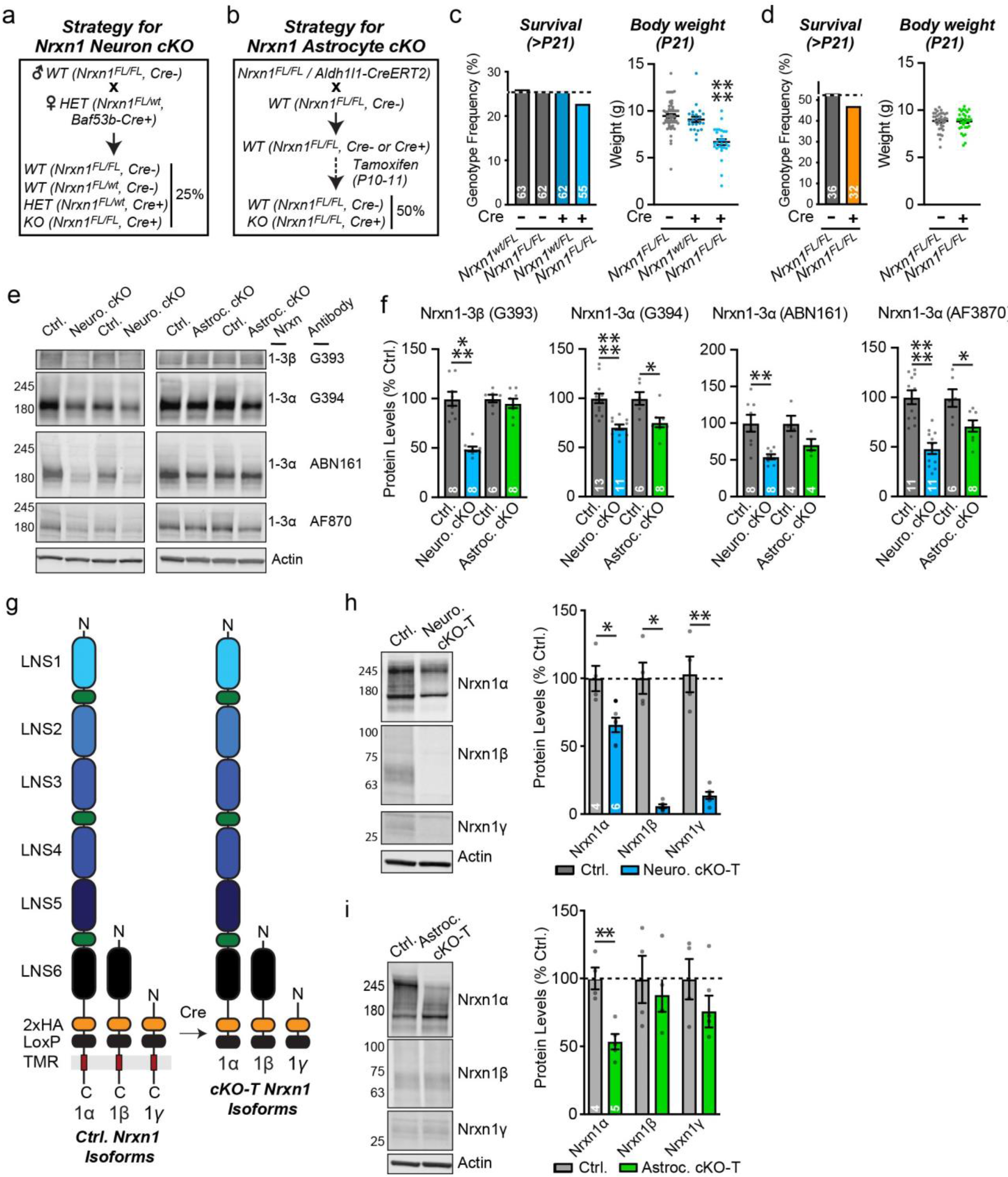
Generation of Cell-Specific Nrxn1 cKO Mice and Quantification of Neurexin Protein Levels (data complement those shown in figures 1-2). **a-b,** Breeding strategies to generate neuron- (**a**), and astrocyte-specific Nrxn1 cKOs (**b**). Note that these breeding strategies operate either with classical Nrxn1 cKO mice or with Nrxn1 conditional HA-knockin mice. All functional studies were performed with classical cKO mice, and conditional HA-knockin mice were only used for studies of the localization and modification of Nrxn1 (Figs. 1 and 2; Extended Figs. 2-6). **c-d,** Effect of neuron- (**c**) and astrocyte-specific Nrxn1 cKOs (**d**) on mouse survival (left graphs) and mouse weight at weaning (right graphs). Numbers in bars represent number of mice examined. e-f, Representative immunoblots of neurexins (**e**) and quantifications of their levels (**f**) using a series of antibodies with different degrees of specificity. Total brain samples from neuron- and astrocyte-specific Nrxn1 cKO mice and littermate controls were examined. Three different antibodies raised against the conserved cytoplasmic domains of Nrxns were used to recognize pan-α-Nrxns (i.e. G394, ABN161, and AF870) and a single antibody was used to recognize pan-β-Nrxn (i.e. G393). Note that most antibodies recognize multiple neurexins, and not only Nrxn1. **g,** Schematic of Nrxn1 proteins produced in conditional Nrxn1 HA-knockin mice. Before Cre-recombination, Nrxn1 HA-knockin mice express HA-tagged Nrxn1; after Cre-recombination, HA-Nrxn1 is truncated and secreted, resulting in cKO-T mice. Because the HA epitope is knocked into the Nrxn1 coding sequence C-terminal to the LNS6 domain followed by loxP sites, Cre-mediated truncated causes production of secreted Nrxn1α, Nrxn1β, and Nrxn1γ fragments that include a C-terminal HA-epitope. These fragments manifest on immunoblots as tight bands that are slightly smaller than wild-type Nrxn1 species (LNS1–6, LNS1-6 domains; E, EGF-like domain). **h-I**, Quantitative immunoblot analysis of neurexins after neuron- (**h**) or astrocyte-specific Cre-recombination (**i**) in Nrxn1 HA-knockin mice creating truncated neurexin variants (left, representative immunoblots using HA-antibodies; right, quantifications of Nrxn1α, Nrxn1β, and Nrxn1γ). Note that owing to the truncated HA-tagged Nrxn1 fragment after Cre-recombination, the deletions do not abolish as much of the Nrxn1 signal as observed in standard Nrxn1 cKO mice analyzed by Nrxn1 antibodies (see Fig. 2). Numerical data are means ± SEM. *n =* indicated on all graphs except for body weight in **c-d** with 60 *Nrxn1^FL/FL^* (cre-), 23 *Nrxn1^wt/FL^* (cre+), and 28 *Nrxn1^FL/FL^* (cre+) mice (**c**, right); 36 *Nrxn1^FL^*^/FL^ (cre-) and 32 *Nrxn1^FL^*^/FL^ (cre+) mice (**d**, right). Statistical significance was assessed with a chi²-test (**c** and **f**, survival), one-way ANOVA with a Tukey’s post-hoc test (**c** and **f**, body weight), and two-tailed unpaired t-test to controls (rest of **f**, **h**, **i**), with * = p < 0.05; ** = p < 0.01; *** = p < 0.001; **** = p < 0.0001.

**Extended Data Figure 5.**
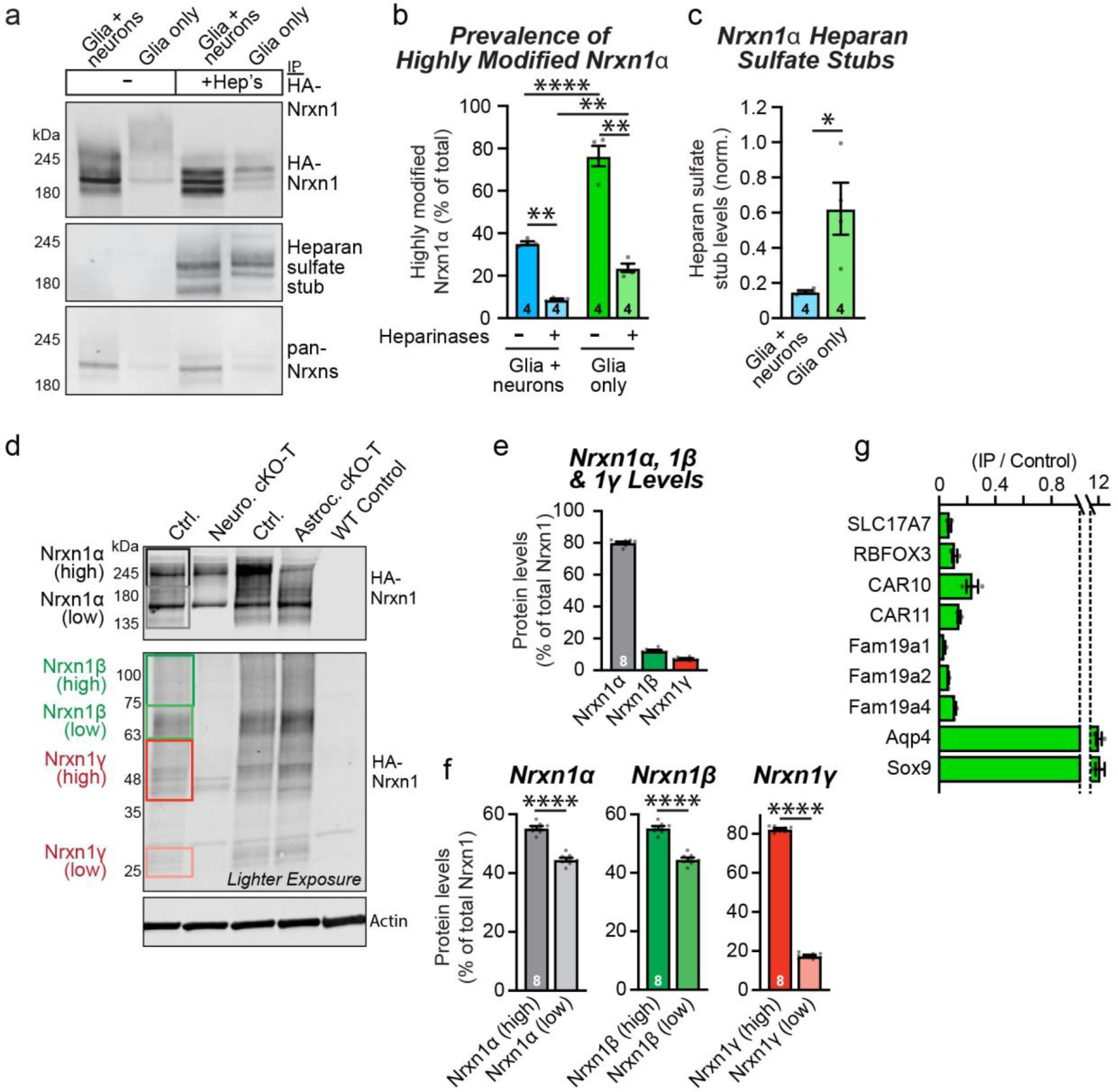
Astrocytic Nrxn1 is Preferentially Modified by Heparan Sulfate (data complement those shown in figure 2). **a-c,** Nrxn1α is expressed primarily as a heparan-sulfate proteoglycan in cultured astrocytes, whereas Nrxn1α is present in cultured neurons mostly without the heparan-sulfate modification. Nrxn1 was immunoprecipitated at DIV14-16 from cultures of glia and neurons, or of glia alone, that were obtained from Nrxn1 HA-knockin mice. Immunoprecipitates were analyzed by immunoblotting without or with treatment with heparinases that cleave heparan-sulfate modifications, exposing a heparan-sulfate stub that can be visualized by an antibody (left, representative immunoblots; right, relative levels of heparan-sulfate stubs normalized for total Nrxn1α levels). Quantification of the amounts of Nrxn1α that are highly modified based on their apparent molecular weight as analyzed by immunoblotting of Nrxn1α immunoprecipitated from cultured glia mixed with neurons or glia alone that were obtained from HA-knockin mice. Note that in the mixed cultures, the relative amounts of highly modified Nrxn1α are decreased relative to the cultures of glia only presumably because more neuronal Nrxn1α is not highly modified. **d-f,** Immunoblot analyses of brain homogenates from conditional Nrxn1 HA-knockin mice, examining control mice and mice with neuron- and astrocyte-specific Cre-expression to create cell type-specific conditional truncations (cKO-T’s). On representative immunoblots (**d**), boxes on the left indicate assignments to specific high- and low-molecular weight neurexin variants. (**e**) Quantification of the total relative amounts of Nrxn1α protein (∼80%), Nrxn1β protein (∼12%) and Nrxn1γ protein (∼7%). (**f**) Quantification of the relative amounts of highly and less modified of Nrxn1α, Nrxn1β, and Nrxn1γ. **g,** Quantification of neurexin-binding, HS modifier transcript enrichment in astrocyte-specific ribosome-bound mRNAs (targeted using RiboTag) compared to total RNA (input). Markers for neurons (i.e. SLC17A7) and astrocytes (i.e. Aqp4 and Sox9) are included. Numerical data are means ± SEM. *n* is indicated on graphs, except 3 pooled biological replicates (2 animals per) in **g.** Statistical significance was assessed with a two-tailed unpaired t-test to controls (**c** and **f**) or two-way ANOVA with a Tukey’s post-hoc test (**b**), with * = p < 0.05; ** = p < 0.01;; **** = p < 0.0001.

**Extended Data Figure 6.**
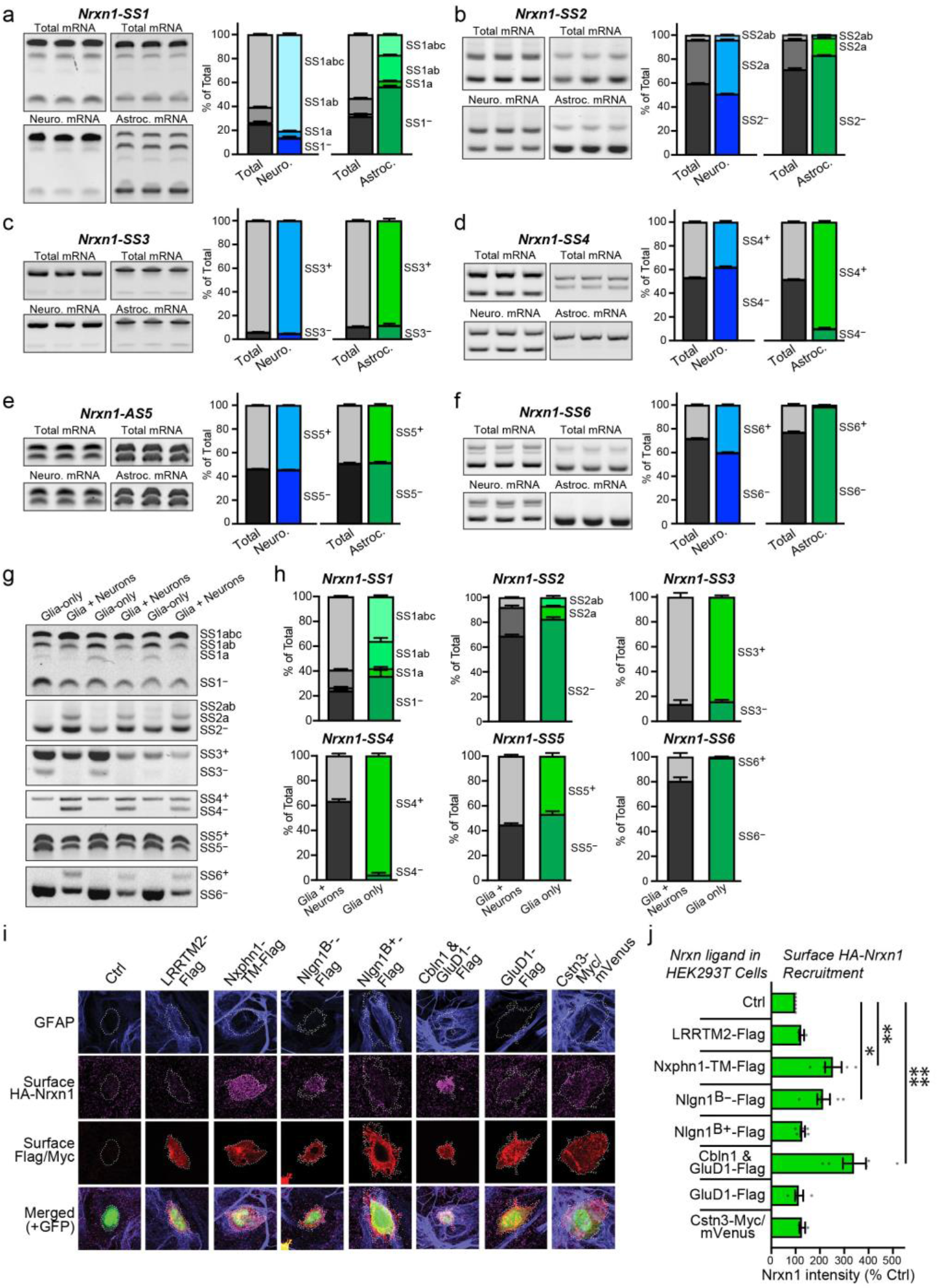
Nrxn1 alternative splicing differs between astrocytes and neurons (data complement those shown in figure 2). **a-f**, Comparison of Nrxn1 alternative splicing at SS1 to SS6 in neurons vs. astrocytes reveals profound differences. Translating mRNAs were immunoprecipitated from neurons or astrocytes using Cre-dependent RiboTag mice crossed with neuron- or astrocyte-specific Cre-driver mice. Alternative splicing was quantified using junction-flanking PCR. Each panel depicts a single site of alternative splicing (left, representative gel images; right, distribution of splice variants in % of total). **g-h**, Comparison of Nrxn1 alternative splicing at SS1 to SS6 in cultures of either glia alone or glia and neurons together confirms profound differences. Junction-flanking PCR was performed on mRNA from DIV14-DIV16 primary hippocampal mixed cultures (with glia and neurons) and pure glia cultures (primarily astrocytes). Representative gel images (**g**) and summary graphs of individual splice sites (**h**). **i-j**, Only a subset of Nrxn1 ligands when expressed in HEK293T cells recruit HA-Nrxn1 expressed by co-cultured astrocytes from HA-Nrxn1 cKI mice. Pure cultures of glia were co-cultured with HEK293T cells expressing the indicated Nrxn1 ligands. After 48 h, surface HA-Nrxn1 recruitment to HEK293T cells was quantified (**i**, representative images; **j**, quantification of HA-Nrxn1 recruitment). Numerical data are means ± SEM. n = 4 mice for neuron mRNAs and 8 mice with 2 mice pooled per pull-down for astrocyte mRNAs (**a-f**); 3 independent cultures (**g-h**); 10-20 cells averaged per culture from 3-6 cultures (**i-j**). Statistical significance was assessed using two-way ANOVA with a Tukey’s post-hoc test (**j**), with * = p < 0.05; ** = p < 0.01;; **** = p < 0.0001.

**Extended Data Figure 7.**
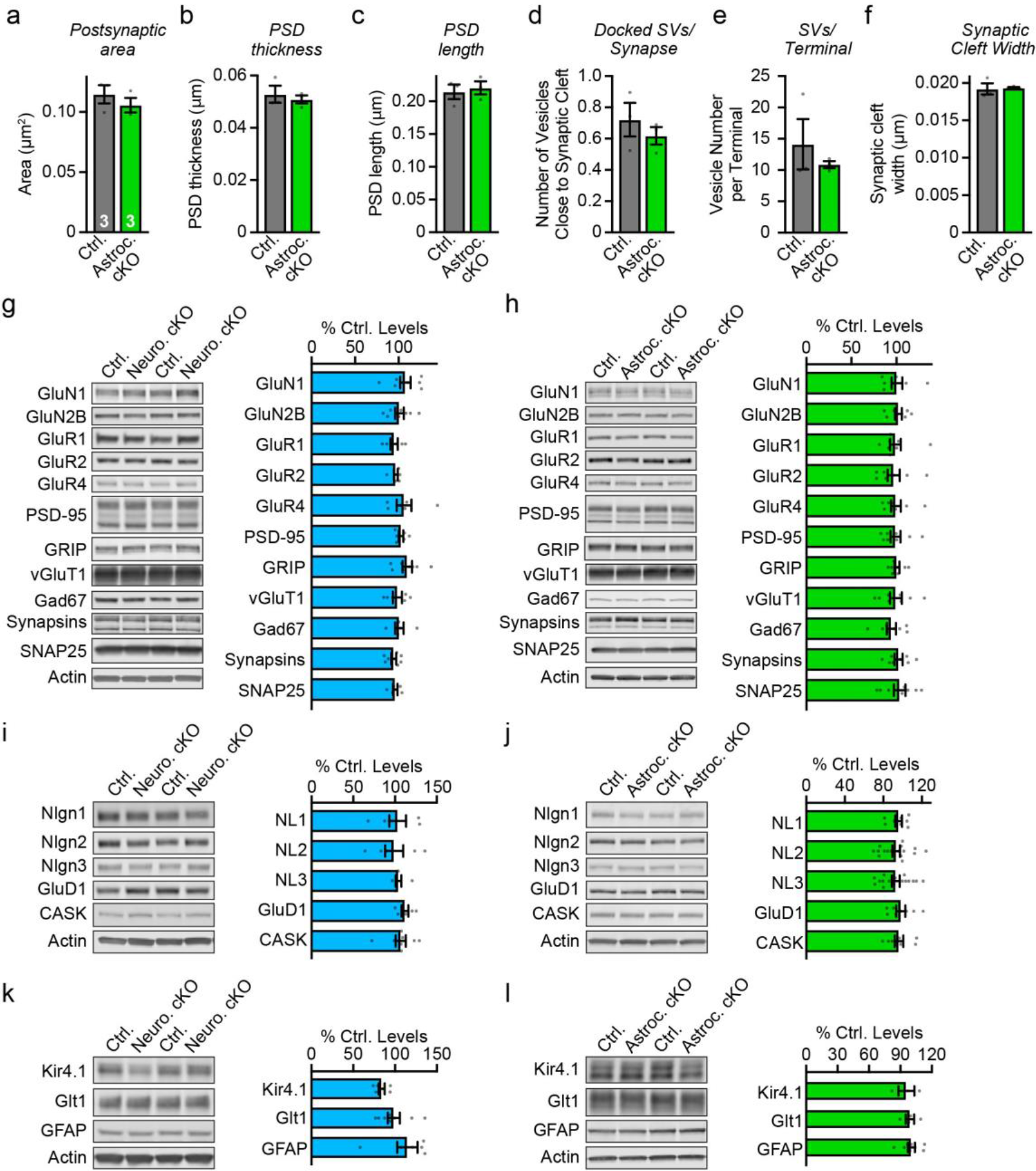
Ultrastructural parameters of excitatory synapses in the CA1 region analyzed in mice with an astrocyte-specific Nrxn1 deletion (a-f) and immunoblotting analyses of the hippocampus of mice with neuron- or astrocyte-specific Nrxn1 deletions (g-l; data complement those shown in figure 3). Ultrastructural data on excitatory synapses after astrocyte-specific deletion of Nrxn1, extending the data shown in Fig. 3g & 3h (**a,** postsynaptic spine area; **b,** PSD thickness; **c,** PSD length; **d,** number of docked synaptic vesicles/synapse; **e,** number of synaptic vesicles per terminal; **f,** width of synaptic cleft). **g-l,** Immunoblotting analyses show that the neuron- (**g, i, k**) or astrocyte-specific (**h, j, l**) Nrxn1 deletions do not produce detectable changes in the levels of selected synaptic proteins (**g, h**), neurexin ligands (**i, j**), or astrocytic proteins (**k, l**) in the hippocampus analyzed at P26-P30. Representative images of immunoblots (left of each column) and summary graphs of protein levels (right of each column; normalized to controls) are shown. Proteins are organized into groups as indicated. Numerical data are means ± SEM. *n* = indicated above except, 6-8 ctrl / 6-8 neuron cKO; 8 ctrl / 8 astrocyte cKO, except Glt1 and Kir4.1 with 4/4 and 3/3, respectively (**g-l**).

**Extended Data Figure 8.**
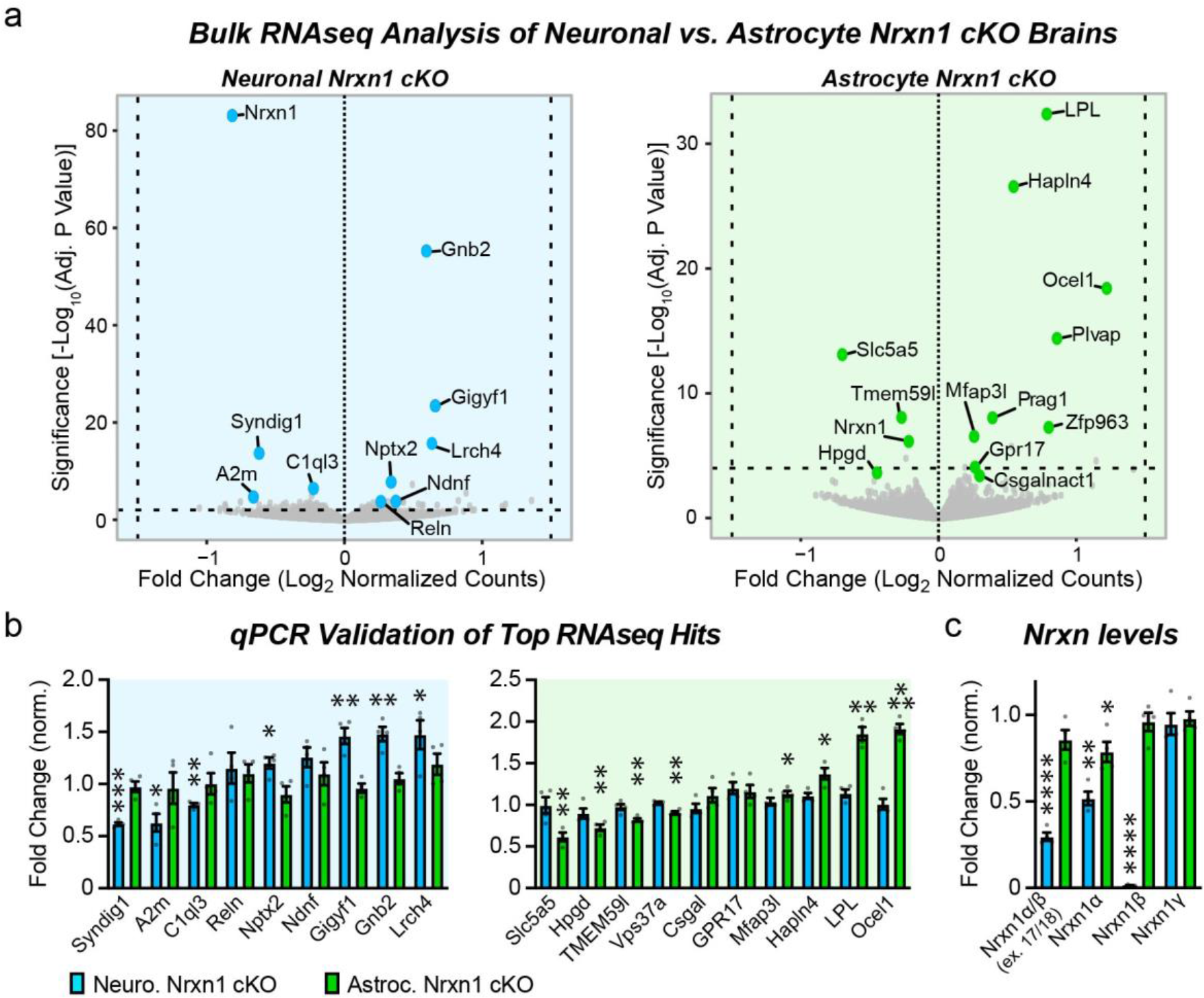
RNAseq reveals that neuronal and astrocytic Nrxn1 control distinct hippocampal transcriptional programs. **a,** Volcano plots of batch-normalized differential gene expression in the hippocampus of neuron- (left) and astrocyte-specific Nrxn1 cKOs (right). Bulk RNAseq analysis was performed at P26 in two batches with 2 control and cKO mice each (2 males and 2 females). **b,** qPCR validation of top differentially expressed genes emerging from RNAseq experiments (left, neuron-specific cKO; right, astrocyte-specific cKO; n = 4 per genotype). **c,** Plot of Nrxn1 mRNA levels as determined by RNAseq validating the quantitative RT-PCR data of Fig. 2. Data in B and C are means ± SEM. Statistical significance was assessed with a two-tailed unpaired t-test to controls (B and C; for statistical analysis of RNAseq data, see Experimental Procedures), with * = p < 0.05; ** = p < 0.01; *** = p < 0.001; **** = p < 0.0001.

**Extended Data Figure 9.**
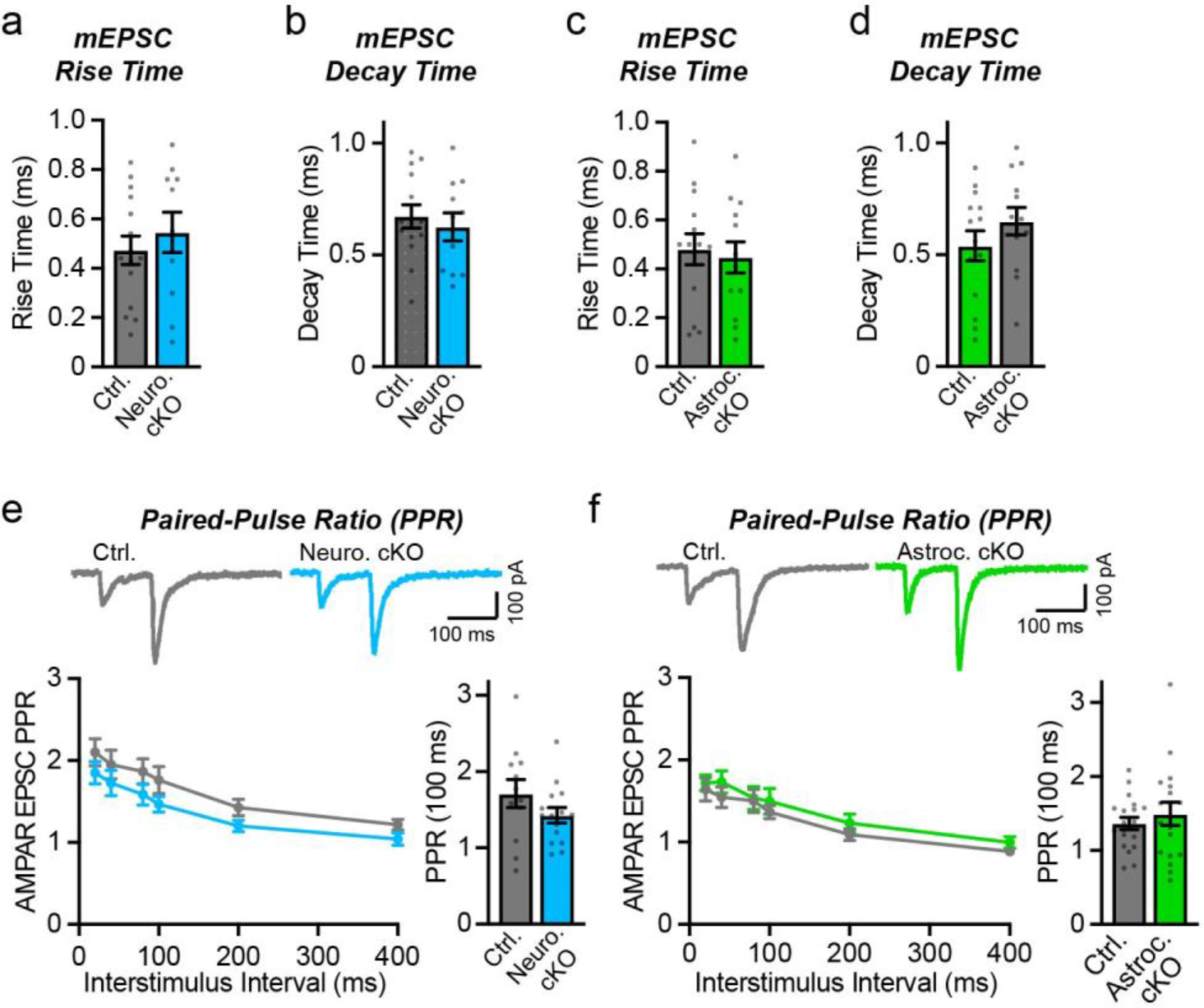
Neuron- or astrocyte-specific deletion of Nrxn1 has no effect on the kinetics of mEPSCs (a-d), and do not alter the paired-pulse ratio of evoked AMPAR-mediated EPSCs (**e, f**). **a-b,** mEPSC rise (**a**) and decay times (**b**) were not significantly altered by the neuron-specific deletion of Nrxn1. (**c-d**) Same as **a-b,** but for the astrocyte-specific deletion of Nrxn1. **e-f,** Neuron- and astrocyte-specific deletions of Nrxn1 do not change the paired-pulse ratios of Schaffer-collateral AMPAR-mediated EPSCs, suggesting that deletion of Nrxn1 in neurons has no effect on release probability (top, sample traces; left bottom, summary plot; right bottom, summary graph of the paired-pulse ratio at a 100 ms interstimulus interval). Numerical data are means ± SEM. *n =* 15 cells / 4 mice ctrl., 11 cells / 4 mice neuron cKO (**a-b**); 14 cells / 4 mice ctrl., 13 cells / 3 mice astrocyte cKO (**c-d**); 12 cells / 3 mice ctrl., 15 cells / 4 mice neuron cKO (**e**); 19 cells / 4 mice ctrl., 18 cells / 4 mice astrocyte cKO (**f**). Statistical significance was assessed with the two-tailed unpaired Student’s t-test.

